# Spatiotemporal single cell transcriptomic analysis of human gut macrophages reveals multiple functional and niche-specific subsets

**DOI:** 10.1101/2021.05.11.443586

**Authors:** Diana Domanska, Umair Majid, Victoria T. Karlsen, Marianne A. Merok, Ann-Christin Beitnes, Sheraz Yaqub, Espen S. Bækkevold, Frode L. Jahnsen

## Abstract

Macrophages are a heterogeneous population of cells involved in tissue homeostasis, inflammation and cancer. Although macrophages are densely distributed throughout the human intestine, our understanding of how gut macrophages maintain tissue homeostasis is limited. Here we show that colonic lamina propria (LpM) and muscularis macrophages (MM) consist of monocyte-like cells that differentiate into multiple transcriptionally distinct subsets. LpM comprise subsets with proinflammatory properties and subsets high antigen presenting and phagocytic capacity. The latter are strategically positioned close to the surface epithelium. Most MM differentiate along two trajectories; one that upregulates genes associated with immune activation and angiogenesis, whereas the other upregulates genes associated with neuronal homeostasis. Importantly, MM are located adjacent to neurons and vessels. Cell-cell interaction and gene network analysis indicated that survival, migration, transcriptional reprogramming, and niche-specific localization of LpM and MM are controlled by an extensive interaction with tissue-resident cells and a few key transcription factors.

## Introduction

Over the last years it has been shown that tissue macrophages, residing in virtually all organs, are extremely heterogeneous. Originally, macrophages were described as professional phagocytes being fundamental to our immune defense. However, it is now clear that macrophages have a broad range of both immune and non-immune functions. Moreover, many macrophage functions are tissue-specific such as electrical conduction in the heart (Hulsmans et al., 2017), synaptic pruning in the brain (Hong et al., 2016), alveolar surfactant clearance in the lung (Guilliams et al., 2013) and limb regeneration in salamanders (Godwin et al., 2013). These functions are thought to be controlled by tissue-specific gene modules regulated by specific transcription factors imprinted by the microenvironment. Interestingly, tissue resident macrophages (TRM) can be reprogrammed when transferred to a new microenvironment (Lavin et al., 2014), indicating that mature TRM retain their plasticity. In addition to the local microenvironment, macrophage functions also depend on their ontogeny (embryonic precursors or adult monocytes), intrinsic factors, and time spent in the tissue (Bleriot et al., 2020). The gut is an organ composed of several compartments with unique functions. The mucosa is covered by a one-cell thick epithelial layer that separates our body from the outside world. This compartment stretches the local immune system to its limits, which has to rapidly and efficiently eliminate pathogens and toxins and at the same time tolerate harmless molecules (e.g. food proteins) and the gut microbiota. Below the mucosa is a thin layer of tissue termed submucosa, which is overlying a thick layer of smooth muscle, termed muscularis propria, consisting of a circular and longitudinal muscular layer. The muscular layers are responsible for segmental contractions and peristaltic movement of the intestinal tract. The gut has an extensive enteric nervous system with a plexus of ganglia cells in both the muscular layers (Auerbach’s plexus) and the submucosa (Meissner’s plexus).

Macrophages populate all layers of the gut wall and studies in mice suggest that they have niche-specific functions that are essential to maintain tissue homeostasis (De Schepper et al., 2018). Under steady state conditions lamina propria macrophages (LpM) constantly phagocytose apoptotic epithelial cells and are critical for gut microbiota composition (Arandjelovic and Ravichandran, 2015; Earley et al., 2018). Recently, LpM in distal colon were shown to maintain epithelial integrity by limiting fungal product adsorption (Chikina et al., 2020). Muscularis propria macrophages (MM) interact with both neurons and vessels and loss of MM leads to loss of enteric neurons, vascular leakage, impaired secretion and reduced intestinal motility (De Schepper et al., 2018; Muller et al., 2014). Moreover, Matheis et al showed that MM protected against post-infectious neuron damage (Matheis et al., 2020). At birth, the mouse gut is populated with fetal-derived macrophages. However, the LpM are rapidly (within weeks) replaced by bone marrow-derived circulating monocytes (Bain et al., 2014). Monocytes also emigrate to muscularis propria, but in this compartment fetal-derived macrophages appear to be more persistent (De Schepper et al., 2018). However, all studies referred to above have been performed in mice, and translation of these results to understand human biology should be made with caution.

Gut macrophages may also be detrimental to the host. Aberrantly activated macrophages have been shown to play a key role in IBD pathology (Martin et al., 2019; Smillie et al., 2019) and tumor-associated macrophages are often associated with worse prognosis (Katzenelenbogen et al., 2020; Zhang et al., 2020). Because of their heterogeneity and plasticity, there is currently a great interest in the search for strategies to reprogram macrophages as a therapeutic tool to treat both inflammatory disorders and cancer (Jahchan et al., 2019). However, to target macrophages in human diseases a deeper understanding of their heterogeneity and tissue-specific functions is necessary.

Our current knowledge about macrophages in the human gut is limited. Using bulk RNA sequencing (RNA-seq) we have shown that LpM the small intestine consist of several transcriptional states (Bujko et al., 2018) and by following their turnover kinetics in transplanted duodenum we found that host circulating monocytes rapidly entered the graft and differentiated into TRM that completely replaced donor LpM within months after transplantation. Here we extend these findings by performing a detailed characterization of both LpM and MM from adult human colon applying single cell (sc) RNA-seq and multi-color immunostaining in situ. Spatiotemporal analysis shows that both compartments contain multiple coexisting TRM subsets with sub-tissular specific functions. We furthermore identify a limited number of transcription factors that may control TRM diversity and we reveal an extensive cross-talk between TRM subsets and tissue resident cells, including stromal cells, epithelial cells, neurons and immune cell lineages. These cell-cell interactions are most likely responsible for the imprinting of tissue-specific macrophage identity.

## Results

### Macrophages in colonic mucosa and muscularis propria comprise transcriptional states associated with monocytes and tissue resident macrophages

Samples from the large intestine were obtained from colorectal cancer resections. Macroscopically and microscopically normal tissue at least 10 cm from the tumor tissue was used. Mucosa and muscularis propria were separated longitudinally and pieces of tissue from each compartment were enzymatically digested to obtain single cell preparations. Cells from mucosa and muscularis were analyzed separately (Figure 1). Flow cytometric analysis revealed that CD45^+^CD3^−^CD19^−^HLA-DR^int/+^ cells separated into either CD14^+^ macrophages or CD14^−^ classical dendritic cells (cDC), including CD141^+^ cDC1 and CD1c^+^ cDC2 (Figure S1A), as previously shown in the small intestine (Bujko et al., 2018). Thus, tissue-derived macrophages were sorted as CD45^+^HLA-DR^+^CD14^+^ (Figure S1B) and processed on a 10X Genomics Chromium platform. A total of 63.970 cells with high quality mRNA were analyzed. To present high dimensional data in low dimension we constructed UMAP plots of each compartment. We found that sorted CD14^+^ cells expressed the macrophage markers CD163 and CD68, further demonstrating that cells included in the analysis were all macrophages (Figure S1C). We have previously shown that the macrophage population in the small intestine contains several transcriptionally distinct subpopulations (Bujko et al., 2018); including “transient” monocyte-like macrophages expressing high levels of calprotectin (heterodimer of S100A8 and S100A9) and long-lived calprotectin-negative macrophages expressing a TRM phenotype (Bujko et al., 2018). To examine whether colonic LpM and MM contained similar phenotypes we first assessed the expression of S100A8 and S100A9, and the complement component 1 chains (C1QA, C1QB, C1QC); the latter highly expressed on small intestinal TRM (Bujko et al., 2018). UMAP visualization of both compartments showed that most LpM and MM expressed either S100A8/S100A9 or C1QA/C1QB/C1QC genes (Figure 2A). Further transcriptomic analysis of S100A8/S100A9^+^ macrophages showed a high number of differentially expressed genes (DEG, e.g. FCN1, VCAN, AQP9, TREM1) that are preferentially expressed in blood monocytes and small intestinal monocyte-like macrophages (Figure 2B). C1QA/C1QB/C1QC^+^ macrophages, on the other hand, expressed many genes associated with TRM (e.g. MRC1, APOE, SELENOP, CSF1R, MERTK, and LYVE1). Flow cytometry of dispersed tissue macrophages confirmed that LpM and MM consisted mainly of calprotein^+^C1q^−^ and calprotectin^−^C1q^+^ macrophages (Figure 2C).

**Figure 1:**
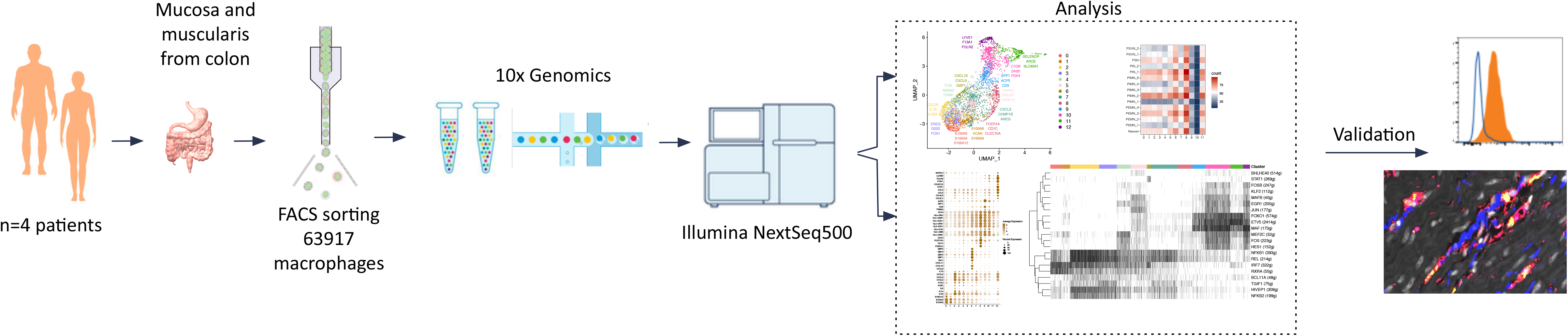
Schematic overview of scRNA-seq analysis of colonic mucosa- and muscularis propia-derived macrophages.

**Figure 2:**
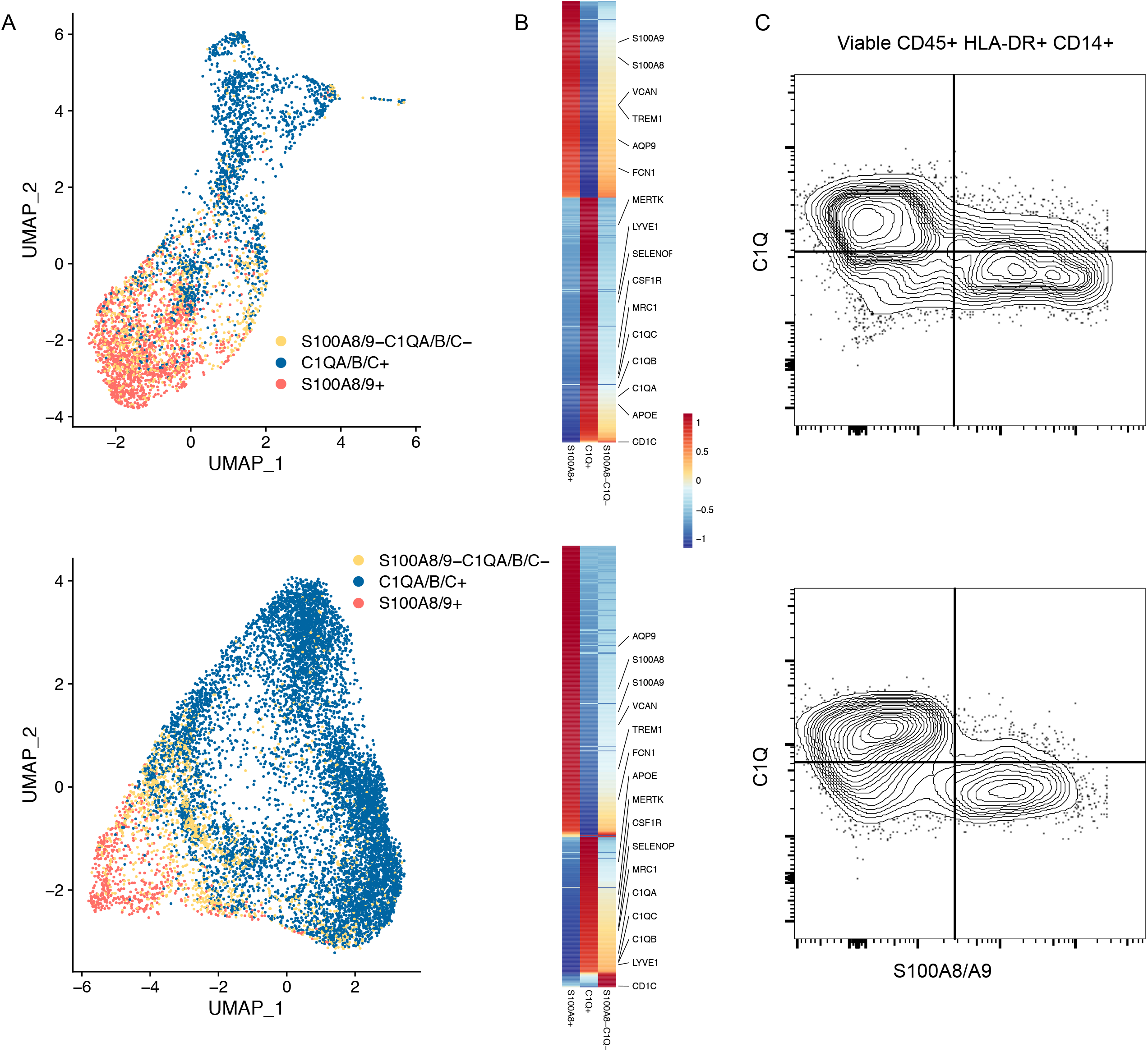
Identification of monocyte-like and tissue-resident macrophages (TRM) in colon mucosa and muscularis propria. A) UMAP plot showing expression of S100A8/A9+ (red), C1QA/B/C+ (blue) and S100A8/A9− C1QA/B/C− (yellow). The color among compartments is determined by its higher expression level. B) Heatmap of differentially expressed genes by S100A8/A9+, C1QA/B/C+ and S100A8/A9− C1QA/B/C− cells. Expression levels are visualized from low expression (blue) to high expression (red). C) Representative flow cytometry plots showing intracellular staining for C1Q and S100A8/S100A9 (calprotectin) in CD45^+^HLA-DR^+^CD14^+^ cells from mucosa (top) and muscularis propia (bottom). Gates were set based on staining with irrelevant isotype-matched antibodies.

Together, these findings demonstrated that both LpM and MM contained monocyte-like macrophages and macrophages with a TRM phenotype.

### Mucosal macrophages comprise transcriptionally distinct and niche-specific subsets

To characterize LpM further we performed clustering analysis using Seurat 3.0 (Stuart et al., 2019). Based on DEG, 13 clusters were identified (Figures 3A and S2) that encompassed macrophages from all patients included in the study (Figure S3A). DEG in clusters 0-4 (LpM_0_-LpM_4_) were reminiscent of small intestinal monocyte-like macrophages (Bujko et al., 2018) and enriched gene ontology (GO) terms were typical for innate immune responses such as responses to bacterium, fungus and LPS (Figure 4). LpM_0_ expressed the highest levels of S100A8, S100A9 and S100A12 (Figure 3B), indicating that this cluster constitutes the most recently elicited monocytes. LpM_2_ expressed high levels of many proinflammatory genes (e.g. IL-1B, IL-1A, IL-6, IL23A, CXCL2, CXCL3, CXCL8) reminiscent of inflammatory macrophages found in IBD lesions (Martin et al., 2019) (Figure 3B, C). The immunoregulatory cytokine IL10 showed highest expression in LpM_2_-LpM_4_ (Figure 3B). LpM_5_ contained DEG encoding many heat shock proteins (Figure S2) and top GO terms were related to unfolded protein responses (Figure 4). Interestingly, LpM_6_ expressed many IFNγ-inducible genes virtually absent in the other clusters; (e.g. CXCL9, CXCL10, CXCL11, IDO1, GBP1, GBP2, GBP4, GBP5) (Figure 3B, C). The top GO terms were antigen processing and presentation of exogenous peptide antigen via MHC class I, and type I IFN-, IFNg- and TNF-signaling pathway (Figure 4). This phenotype has recently been reported to promote anti-tumor immunity in colorectal cancer of mice (Qu et al., 2020). Cluster 8 expressed low levels of both S100A8/S100A9 and C1Q genes (Figure 1), but many DEG were associated with DC (e.g. FCER1A, CD1C, CLEC10A, CD1E) (Figure 3B). This cluster also expressed high levels of MHC class II genes (Figure 3B, C) and top GO term was antigen processing and presentation of exogenous peptide antigen via MHC class II (Figure 4). Flow cytometric analysis confirmed the presence of CD14^+^FcER1^+^CD1c^+^ cells in dispersed mucosa (Figure S4). This phenotype is reminiscent of a recently identified DC population - termed DC3, which originates independently of both classical DC and monocytes (Bourdely et al., 2020).

**Figure 3:**
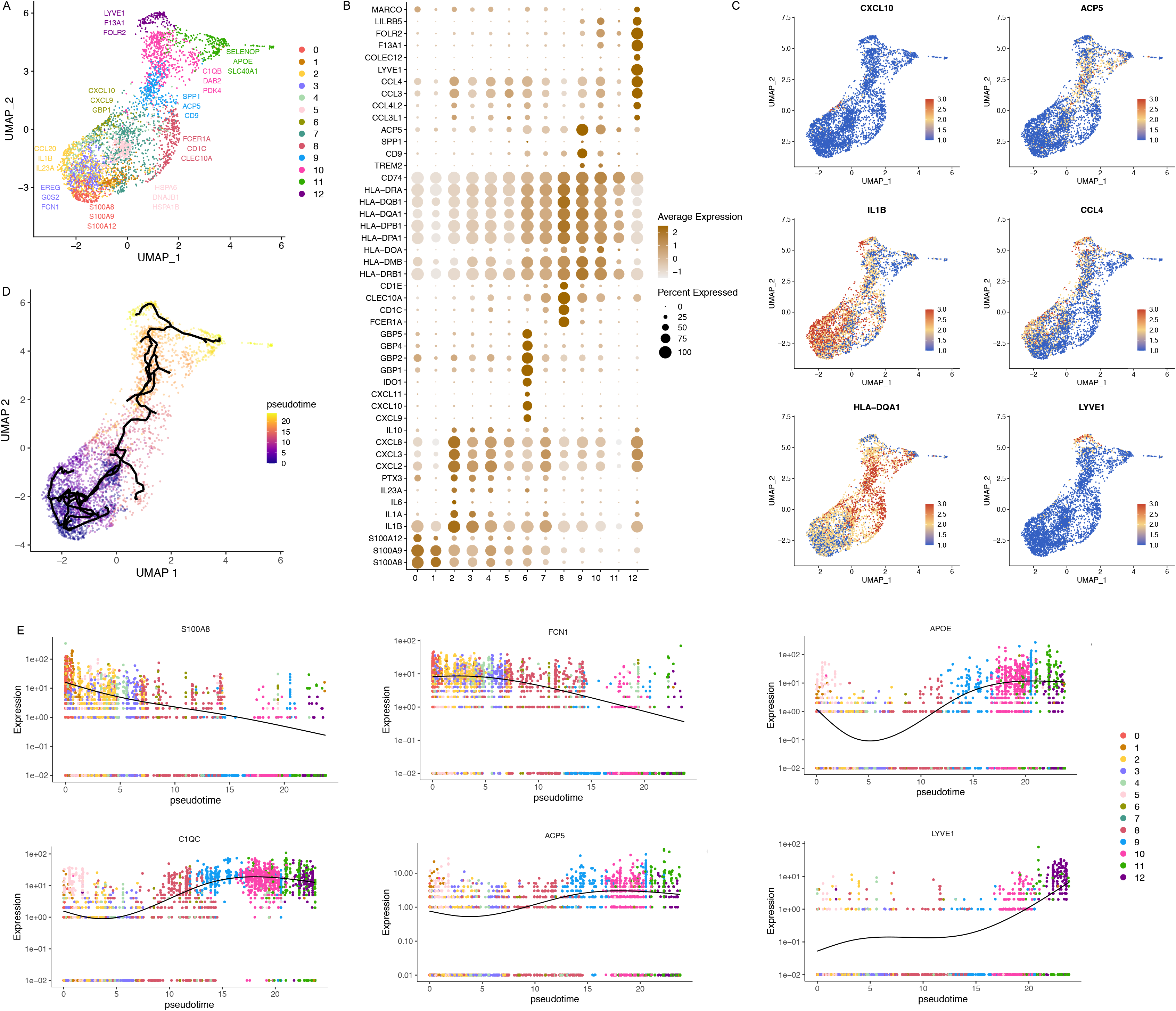
Cluster analysis, differential gene expression identification and pseudotime trajectories for LpM. A) UMAP representation of 13 clusters (different colors) with annotated highly expressed gene identity with in selected clusters. B) Dot plot of average expression of jselected genes. Expression levels are visualized from low (white) to high expression (brown). The percentage of positive cells is indicated by circle size. C) UMAP visualization of selected genes. Expression levels are visualized from low (blue) to high expression (red). D) Reconstructed developmental trajectory of LpM showing pseudotime colored from blue to yellow. E) Expression for some selected gene expression through pseudotime. Cell colors represents clusters.

**Figure 4:**
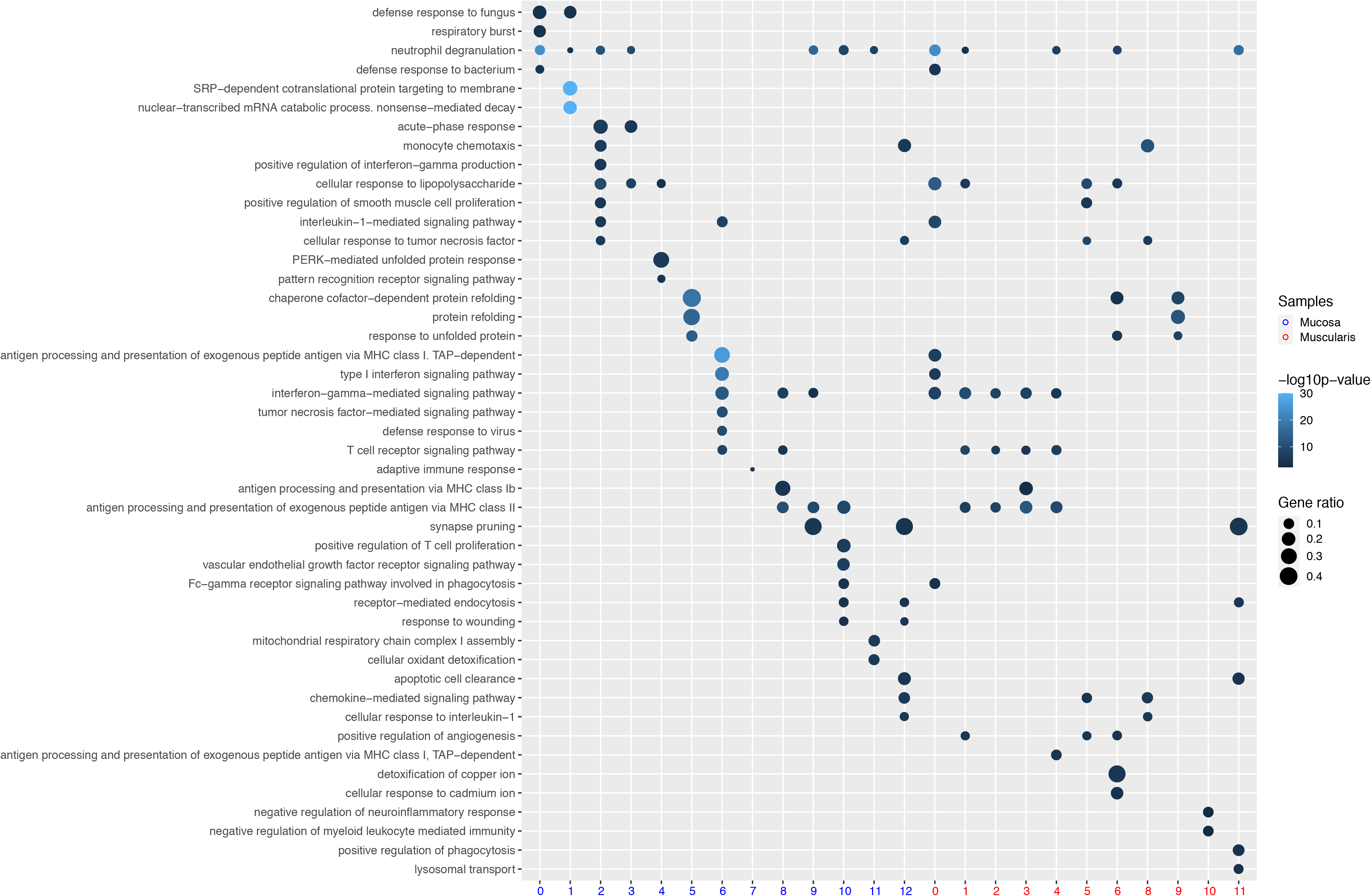
Gene ontology (GO) terms significantly (p< 0.05) enriched at the transcriptomic level for selected clusters of LpM and MM. Gene ratio is represented by circle size.

LpM_9_ and LpM_10_ expressed high levels of HLA class II genes (Figures 3B, C). Accordingly, enriched GO terms were antigen processing and presentation of exogenous peptide antigen via MHC class II (Figure 4). GO terms for LpM_10_ also included wound healing and receptor-mediated endocytosis (Figure 4). Interestingly, LpM_9_ (and to a lesser extent LpM_10_) expressed CD9, TREM2, SPP1, and ACP5 (Figure 3B, C), genes that were recently shown to identify monocyte-derived macrophages enriched in liver fibrosis (Ramachandran et al., 2019) and in adipose tissues of obese patients (Jaitin et al., 2019). LpM_12_ selectively expressed many typical TRM genes, such as LYVE1, COLEC12, F13A1, and FOLR2 (Figure 3B, C), and several chemokine genes (e.g. CXCL2, CXCL3, CXCL8, CCL3, CCL4, (Figure 3B, C). GO terms included chemokine-mediated signaling pathway, synapse pruning, apoptotic cell clearance and receptor-mediated endocytosis (Figure 4).

In agreement with other reports (Bain et al., 2014; Bujko et al., 2018) our findings strongly suggested that the vast majority of LpM originates from bone marrow-derived monocytes, which constantly emigrate to the mucosa and differentiate into TRM. To further understand the process of monocyte-to-macrophage differentiation we reconstructed their developmental trajectories using the Monocle 3 algorithm (Trapnell et al., 2014). We excluded DC3 from the analysis because this subset is reported to originate independently of the monocyte lineage (Bourdely et al., 2020). Selecting random cells in LpM_0_ as root, we identified at least two distinct branches. A short branch ended in Cluster 2, compatible with a distinct proinflammatory trajectory, whereas the majority of the cells went through a more extensive reconfiguration program from LpM_0_ to LpM_9-12_ as a function of pseudotime (Figure 3D). As expected, genes typical for monocytes were downregulated following this trajectory, whereas typical TRM genes were increased (Figure 3E). Together, the findings clearly showed that LpM consisted of multiple transcriptional cell states indicating that incoming monocytes constantly differentiate into multiple distinct TRM subsets with time (Bujko et al., 2018).

Studies in mice have shown that TRM occupy distinct tissue niches where they display niche-specific functions (Chakarov et al., 2019; Guilliams et al., 2020). To determine the anatomical localization of LpM subsets we performed multi-color immunofluorescence stainings in situ. Calprotectin-expressing LpM were found scattered in the lower part of the mucosa between crypts (Figure 5A), whereas C1q was strongly expressed by LpM positioned in the subepithelial region (Figure 5B). To further determine the localization of distinct TRM clusters we stained for Acp5 (expressed by LpM_9_ and LpM_10_), Lyve-1 and Colec12 (expressed by LpM_12_). Interestingly, whereas Acp5^+^ LpM were located in the subepithelial region (Figure 5C), Lyve-1^+^ and Colec12^+^ macrophages were confined to the submucosal region (Figures 5D and S5A) and not found in the mucosa. The latter finding indicated that LpM_12_ represented submucosal macrophages (SmM). Several studies have shown that Lyve-1 is associated with perivascular macrophages, whereas macrophages expressing Colec12, a scavenger receptor for uptake of myelin, has been associated with neurons (Bogie et al., 2017). Here we found that the vast majority of SmM expressed both Lyve-1 and Colec12.

**Figure 5:**
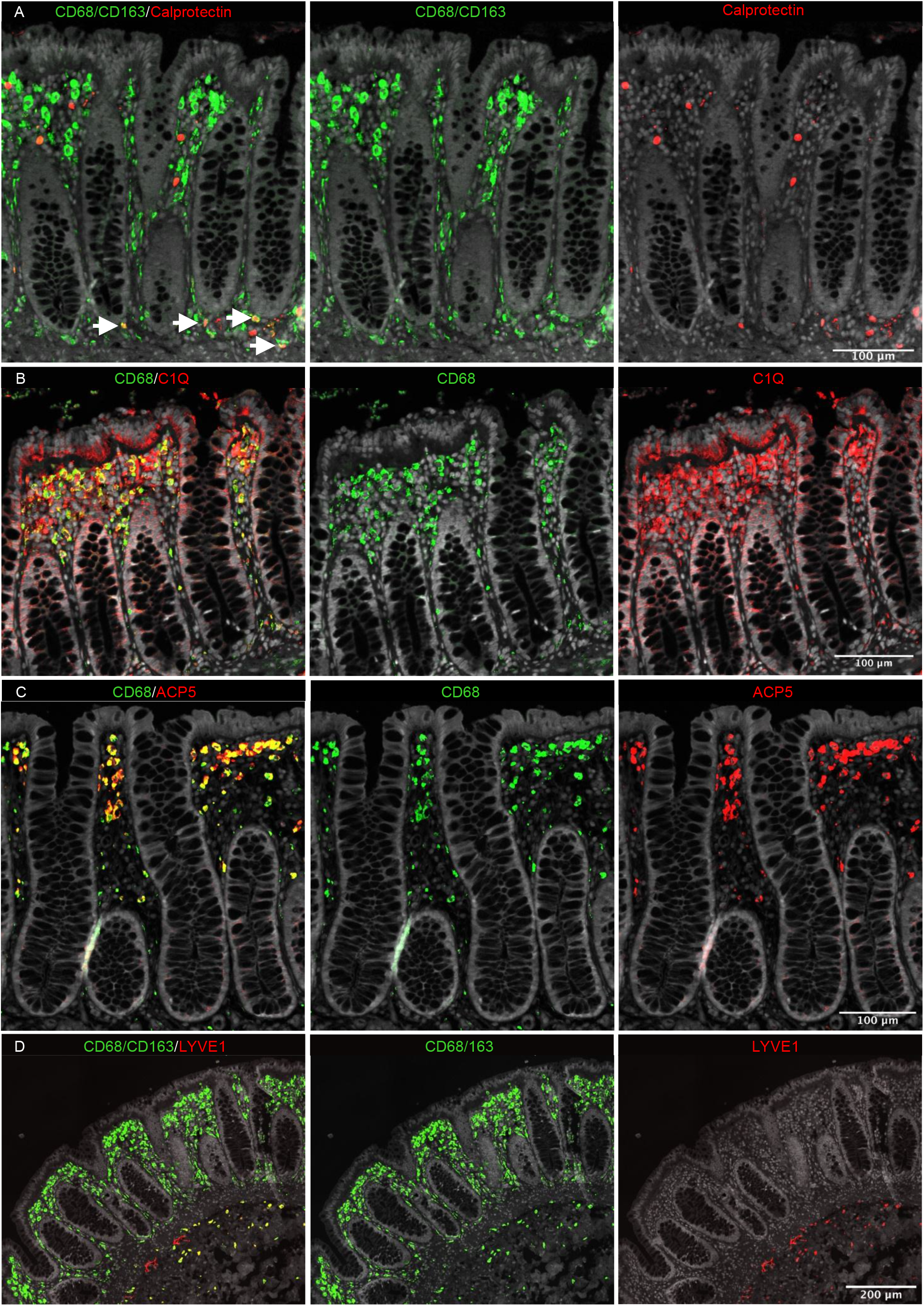
In situ localization of LpM in human colon. Sections stained for A) CD68/CD163 (green) and calprotectin (S100A8/S100A9; red); B) CD68 (green) and C1Q (red); C) CD68 (green) and Acp5 (red); D) CD68 and CD163 (green) and Lyve-1 (red). All sections were counterstained with Hoechst DNA-stain (gray). Arrows indicate CD68/CD163^+^ calprotectin^+^ LpM (yellow) (A). Calprotectin^+^ CD68/CD163^−^ cells (red, left upper panel) are granulocytes. Representative of n≥3.

Collectively, our analysis indicated that the colonic mucosa contains monocyte-derived LpM that differentiate into multiple LpM subpopulations; some subsets with proinflammatory properties (e.g. LpM_2_ and LpM_6_), and other subsets with high antigen presenting and phagocytic capacity that were strategically positioned in the subepithelial region (LpM_9_ and LpM_10_). In contrast, LYVE1^+^ SmM displayed a transcriptomic profile indicating low antigen presenting capacity, but with high chemotactic and tissue-protective properties.

### Muscularis propria macrophages comprise transcriptionally distinct subsets displaying different developmental trajectories

Next, we analyzed macrophages isolated from the muscular compartment. Following high-resolution clustering, the cells were separated into 12 transcriptionally distinct clusters (Figures 6A and S6) encompassing cells from all donors (Figure S3B). In three of the patients, we sampled muscularis propria from two different sites. Clustering analysis revealed that MM from these sites were very similar (Figure S7) indicating that MM subpopulations were distributed homogeneously throughout the muscular compartment. Cluster 0 (MM_0_) expressed high levels of S100A8, S100A9, S100A12, IL1B, IL1A and CXCL chemokines (Figure 6B, C), and enriched GO terms were cellular response to bacterium and LPS as well as type I- and IFNγ-mediated signaling pathway together with MHC class I-mediated antigen presentation (Figure 4). MM_1_, MM_2_, and MM_4_, which clustered adjacent to MM_0_, expressed genes associated with immune activation (e.g. HLA class II genes) (Figure 6B, C) and enriched GO terms were IFNg-mediated signaling pathway and antigen processing and presentation of exogenous peptide antigen via MHC class II (Figure 4). Cluster 3 was reminiscent of DC3 (Bourdely et al., 2020), demonstrating that this DC subset also resided in muscularis propria (Figure S4). The clusters transcriptionally most distant from MM_0_, could be broadly divided into clusters with “proinflammatory” and “homeostatic” properties, respectively. MM_5_ (and MM_8_) expressed high levels of proinflammatory cytokines (e.g. IL1A and IL1B) and multiple chemokines (e.g. CXCL2, CXCL3, CXCL8, CCL3, CCL4), whereas MM_11_ expressed low levels of proinflammatory cytokines and chemokines, but high levels of genes such as LILRB5, MARCO, LYVE1, FOLR2 and COLEC12 (Fig 6B, C). Consistently, enriched GO terms for “proinflammatory” MM_5_ were cellular response to LPS and chemokine-mediated signaling pathway, whereas top GO terms for “homeostatic” MM_11_ were receptor-mediated endocytosis, synaptic pruning and apoptotic cell clearance (Figure 4). Interestingly, high expression of PMP22 and EMP1 genes were observed in MM_8_ and MM_11_ (Figures S6 and S8). These genes are mainly expressed in Schwann cells (Taylor et al., 1995). PMP22 protein is part of the myelin sheath that protects neurons. High expression of PMP22 and EMP1 genes suggests that MM are phagocytosing Schwann cells and thus are in intimate contact with enteric neurons. Cluster 9, similar to LpM_8_ in mucosa, expressed high levels of heat shock protein genes (HSP) (Figure S6) with enriched GO terms such as unfolded protein responses (Figure 4). HSP protects against cellular stress and it was recently shown that increased expression of HSP in macrophages concomitant with downregulation of IL-1 had anti-inflammatory effects in response to change of diet in experimental mice (Brykczynska et al., 2020). In agreement, HSP+ MM expressed low levels of proinflammatory cytokines and chemokines (Figure 6B).

**Figure 6:**
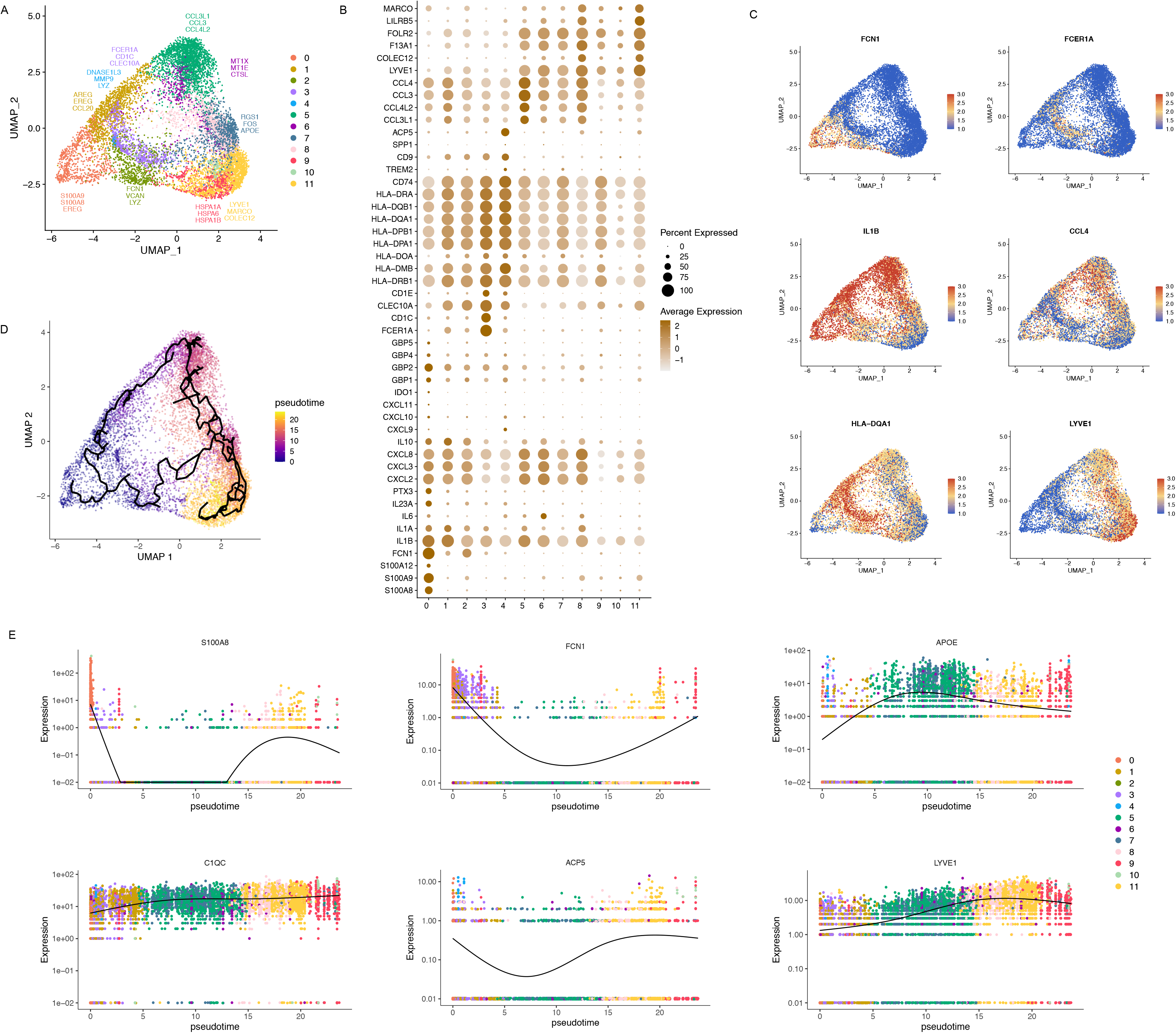
Cluster analysis, gene expression identification and pseudotime trajectories for MM. A) UMAP representation of 12 clusters (different colors) with highly expressed genes within selected clusters. B) Dot plot of average expression of selected genes in every cluster. Average expression levels are visualized from low expression (white) to high expression (brown). The percentage of positive cells is indicated by circle size. C) UMAP visualization of selected genes. Expression levels are visualized from low expression (blue) to high expression (red). D) Pseudotime trajectory analysis of MM. Cells are labeled by pseudotime colored from blue to yellow. E) Expression for some selected gene expression through pseudotime. Cell colors represents clusters.

**Figure 7:**
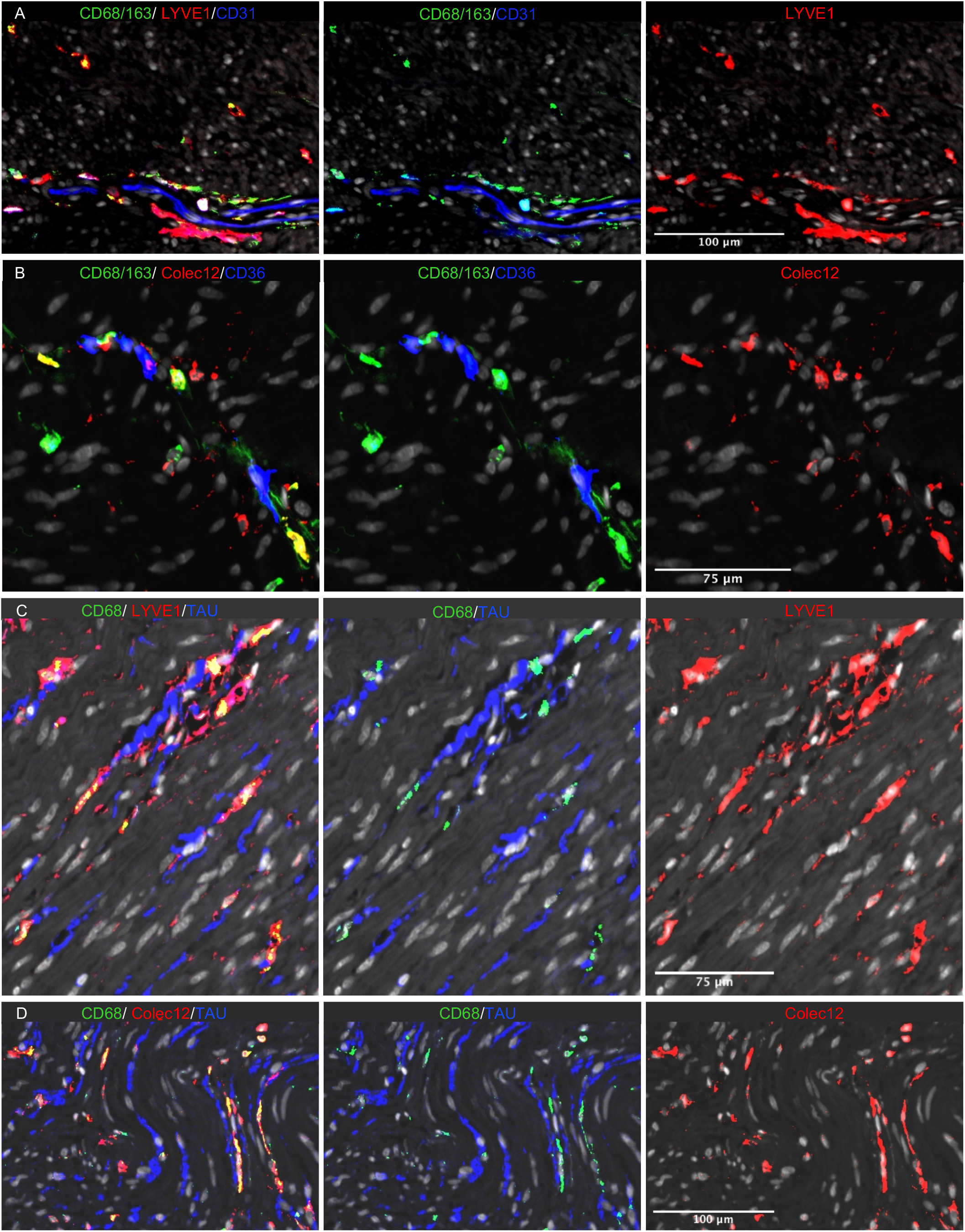
In situ localization of MM in human colon. Sections stained for A) CD68/CD163 (green), LYVE1 (red), and endothelial cell marker CD31 (blue); B) CD68/CD163 (green), Colec12 (red), and endothelial cell marker CD36 (blue); C) CD68 (green), neuron marker tau (blue), and lyve-1 (red); D) CD68 (green), tau (blue), and Colec12 (red). All sections were counterstained with Hoechst DNA-stain (gray). Representative of n≥3.

Pseudotime trajectory analysis, using random cells in MM_0_ as root, showed a trajectory following two distinct branches (Figure 6D). One branch followed through MM_1_, MM_5_, and MM_6_; clusters expressing many proinflammatory genes. Interestingly, all three subsets shared the GO term positive regulation of angiogenesis (Figure 4). Conversely, “homeostatic” branch expressed lower levels of proinflammatory genes (Figure 6B) and ended in MM_11_; the cluster most strongly associated with enteric neurons. As for LpM, typical monocyte-related genes were rapidly downregulated along the trajectory, whereas TRM genes (e.g. LYVE1, C1QC and APOE) were rapidly upregulated and found to be more broadly expressed by MM (Figure 6E) than by LpM (Figure 2E). To determine the anatomical localization of MM we performed multicolor immunostaining in situ. As in the submucosa, most MM expressed both Lyve-1 and Colec12.

They were distributed throughout the muscularis tissue but were enriched adjacent to neurons and vessels (Figure 6A-D). In addition, a minor fraction of calprotectin^+^ MM were found scattered throughout the tissue (Figure S5B).

Together, we found that the MM population was very heterogeneous, consisting of multiple functionally distinct subsets. Transcriptomic profiling and pseudotime trajectory analysis revealed significant differences between MM and their LpM counterparts, indicating that tissue-specific signals from the local microenvironment are important for macrophage differentiation and diversity.

### A subpopulation of mucosal and muscularis macrophages express genes compatible with an embryonic origin

Studies in mice have reported that gut macrophages are composed of both monocyte-derived and embryonic-derived macrophages (De Schepper et al., 2018; Shaw et al., 2018). To analyze whether embryonic-derived macrophages also occurred in adult human colon we examined genes reported to be differentially expressed between lineages. Interestingly, MM_4_ and LpM_9_ (and MM_10_) showed higher expression of several genes related to embryonic ontogeny, such as CD63, ADAMDEC1, and DNASE1L3 (Figure S9A, B) (De Schepper et al., 2018; Shaw et al., 2018). Moreover, using ClusterMap (Gao et al., 2019), a method to determine cluster similarity across biological samples, we found that MM_4_ and LpM_9_ showed the highest similarity index (0.21) when all clusters were compared to each other. To further investigate the possibility that these clusters contained embryonic-derived macrophages we examined the transcriptomic profile of intestinal macrophages in human embryos. Fawkner-Corbett et al. recently published a study analyzing the human intestinal development using scRNA-seq (Fawkner-Corbett et al., 2021). Reexamining the immune cell data of embryos at post conception week 12-22 we found that intestinal macrophages contained three distinct subpopulations (Figure S9C). Cluster 3 expressed high levels of monocyte-related genes such as S100A8, S100A9, VCAN, and FCN1, whereas cluster 0 and 4 expressed higher levels of genes associated with TRM such as C1QA and SELENOP (Figure 9D). Interestingly, only cluster 4 expressed high levels of DNASE1L3 and ADAMDEC1, similar to embryonic-derived macrophages in mouse colon (Figure 9D). Together, these results may suggest that a minor population of gut macrophages in human adults are of embryonic ontogeny.

### Mucosal and muscularis macrophages interact extensively with resident tissue cells and immune cells

Cell differentiation in tissues is triggered by contacts with other neighboring cells through receptor-ligand interactions. Thus, we interrogated microenvironmental signals that could be involved in the transcriptional reconfiguration process observed in both compartments. We determined such interactions applying CellPhoneDB2.0 (Efremova et al., 2020) combining our scRNA-seq datasets of LpM and a published scRNA-seq dataset covering all stromal and immune cells from normal colonic mucosa (Smillie et al., 2019). Numerous statistically significant interactions were found between LpM and subtypes of fibroblasts, endothelial cells and epithelial cells (Figure 8A). Among receptor-ligand pairs that are particularly important for macrophage survival and differentiation we found that the CSF1R-ligand cytokines CSF1, CSF3 and IL34 (Lavin et al., 2015), were expressed by postcapillary venules and activated fibroblasts, whereas genes involved in the Notch signalling pathway (e.g. DLL4/JAG1:NOTCH2) were expressed by epithelial and stromal cells (Figure S10). Endothelial cells and fibroblasts expressed many adhesion molecule and chemokine genes that interacted primarily with LpM_0_-LpM_8_. This is in agreement with the concept that endothelial cells and fibroblasts are involved in the recruitment, migration and localization of LpM (Figure S10). LpM_9_ and LpM_10_, on the other hand, displayed interactions with epithelial cells (CDH1:aEb7, DSG2:DSC2) (Figure S10) consistent with their subepithelial localization (Figure 5C). Importantly, stromal and epithelial cells showed several interactions with LpM that regulate macrophage functions. This included interactions associated with negative regulation of macrophages activation (LGALS9:HAVCR2, HLA−G:LILRB2, HLA−G:LILRB1, TNFSF10:RIPK1) (Chen et al., 2018; Hartwig et al., 2017; Ocana-Guzman et al., 2016), M2-like polarization (GAS6/PROS1:AXL, GAS6:MERTK) (Myers et al., 2019), and “don’t eat me” signals (CD47:SIRPA, CD52:SIGLEC10) (Barkal et al., 2019; Li et al., 2021). On the other hand, LpM expressed ligands (EGFR and ERRB3) for receptor tyrosine kinases expressed by epithelial cells and activated fibroblasts, suggesting a bidirectional regulation of survival and function between LpM and tissue resident cells. LpM also showed numerous interactions with immune cell subtypes (Figure 8B). In particular, LpM_6_, LpM_7_, LpM_9_, (as well as DC3-like LpM_8_), which expressed high levels of MCH class II genes (Figure 3B), showed interactions with cycling, memory and regulatory T cells (e.g. CD28:CD80, CLTA4:CD80, CD28:CD86, CLTA4:CD86) (Figure S11), suggesting that LpM subsets play a role as regulators of T cell responses in the colonic mucosa. Moreover, selective expression of several CXCR3- binding chemokines were expressed by LpM_6_ (Figures 3B and S11), indicating a role in the recruitment of CD8 T cells.

**Figure 8:**
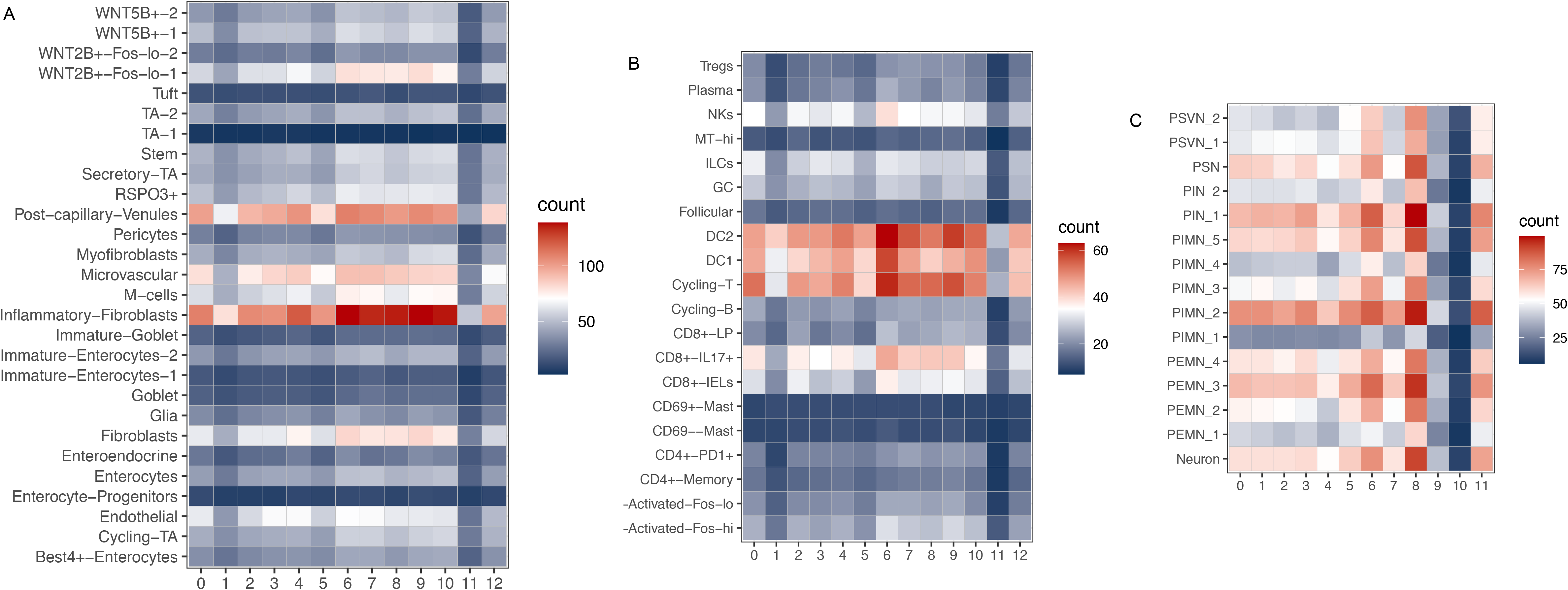
Cell-cell interaction analysis. A, B) Heat map representing numbers of receptor-ligand pairs between LpM clusters and subtypes of epithelial and stromal cells (A), immune cell lineages (B) and subtypes of enteric neuronal cells (C) from normal colon. Rows represents ligand-receptor pairs and columns defines cell-cell interaction pairs.

Studies in mice have shown that MM crosstalk with enteric neurons (De Schepper et al., 2018; Muller et al., 2014). To interrogate macrophage-neuron interactions in muscularis propria we analyzed our MM scRNA-seq dataset together with a published scRNA-seq dataset of the human enteric neuron system (Drokhlyansky et al., 2020). A very high number of interactions between multiple neuron subtypes and MM was found (Figure 8C). Central to MM survival and differentiation, several neuron subtypes expressed Notch ligands (DDL1, DDL3, and JAG2) and IL34 interacting NOTCH2 and CSF1R on MM, respectively. Furthermore, numerous receptor-ligand pairs were involved in macrophage migration, localization and activation/regulation (Figure S12). Finally, MM-neuron interactions were associated with synapse pruning (C3:C3AR1, C5:C5AR2) (Hong et al., 2016) and neuron stimulation (BMP2:BMPR2), strengthening the notion that there is an extensive crosstalk between MM and enteric neurons in the human colon that controls gastrointestinal motility (De Schepper et al., 2018; Muller et al., 2014).

Together, our results indicated that macrophages interact extensively with tissue resident cells in both compartments, compatible with the concept that the local microenvironment is important for imprinting of macrophage specialization and niche-specific localization.

### Gene regulatory network analysis indicates that a limited number of transcription factors control macrophage differentiation and diversity

To identify transcriptional factors (TF) that may control transcriptional programming of LpM and MM we used single-cell regulatory network inference and clustering (SCENIC) (Van de Sande et al., 2020) to determine sets of genes co-expressed with their associated TF (regulons). We found a total of 185 and 187 regulons (active in more than 1% of the cells) for LpM and MM, respectively, with significantly enriched motifs for the corresponding TF. By ordering macrophage clusters along the pseudotime developmental trajectory, we identified a restricted number of regulons corresponding to specific clusters. LpM_0_ showed increased regulon activity corresponding to RXRA and IRF7, whereas LpM_2_ was associated with several NFKB family members (NFKB1, NFKB2, and REL) (Figure 9A). LpM_6_ was associated with STAT1, whereas LpM_8_-LpM_11_ and LYVE1+ SmM showed regulon activity driven by TF such as MAF, MAFB, HES1, and EGR1. MAF and MAFB are known for their role in driving terminal macrophage differentiation (Aziz et al., 2009).

**Figure 9:**
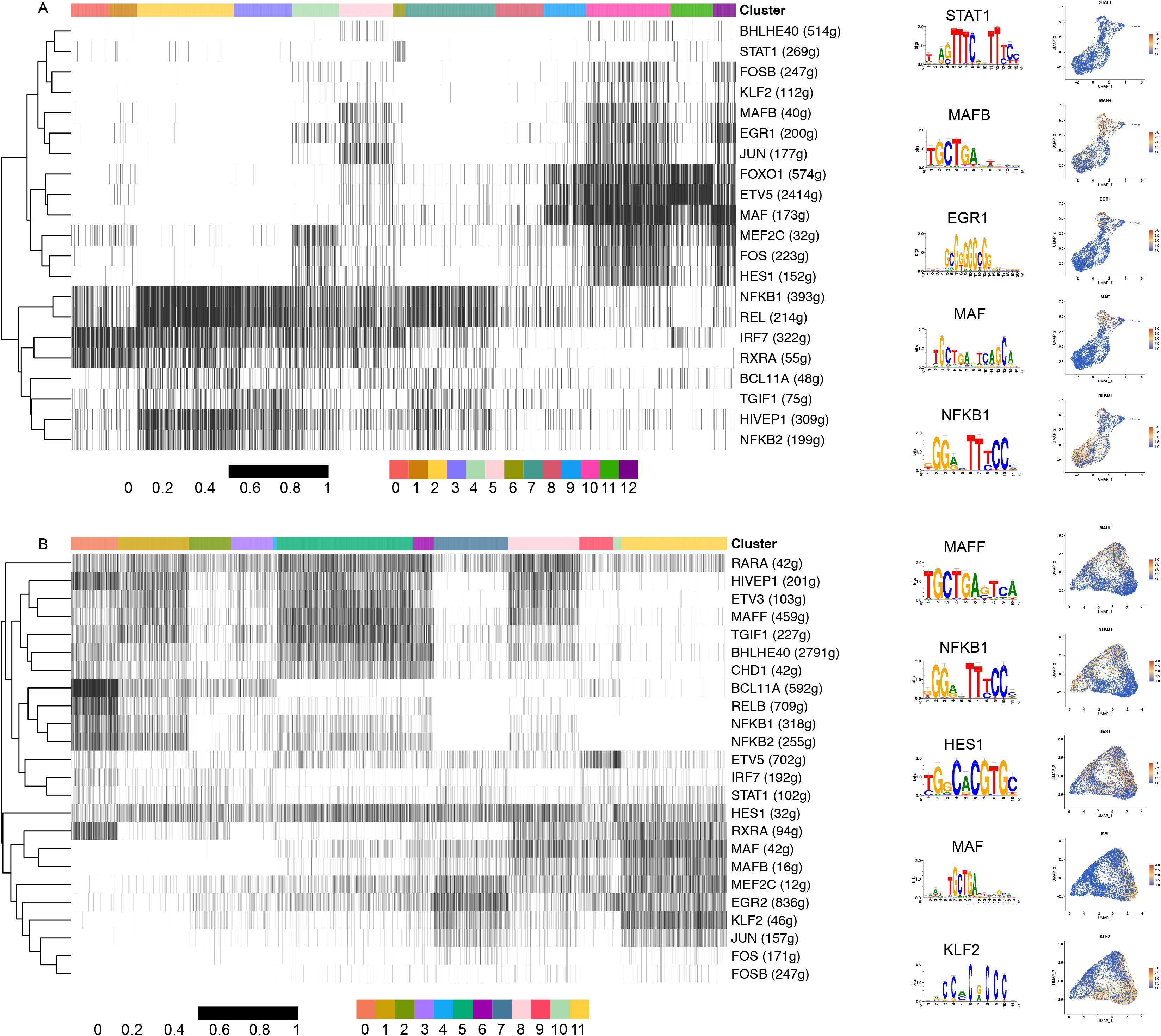
Identification of transcription factor modules based on regulon activities in LpM and MM. Binarized heatmaps of regulon activity in LpM (A) and MM (B) with the representative TF and part of their binding motifs. Numbers in parenthesis indicate the number of genes enriched in the regulons, and UMAP plots show expression of selected TF.

The gene regulatory network in MM showed similarities with LpM (Figure 9B). MM_0_ showed regulon activity corresponding to NFKB family members (NFKB1, RELB and BCL11A), whereas regulons corresponding to MAF and MAFB were upregulated in MM_11_. On the other hand, MM_5_ (and to a lesser extent MM_8_) expressed regulons corresponding to TF such as MAFF, RARA, and BHLHE40, not found in the mucosal compartment. Interestingly, regulon activity with corresponding HES1 was broadly expressed by MM and by subsets of LpM (Figure 9A, B). HES1 is a classic TF downstream of the Notch signalling pathway (Sharma et al., 2012). Together with our cell-cell interaction data (Figures S10 and S12), expression of HES1 suggests that the Notch signalling pathway is involved in reprogramming of LpM and MM, as shown for monocyte-derived macrophages in the liver (Ramachandran et al., 2019; Sharma et al., 2020).

In summary, we find that regulon activity corresponds to distinct macrophage clusters following the pseudotime trajectory, indicating that a limited number of key TF control the process of macrophage reprogramming observed in both compartments.

## Discussion

Using high-resolution spatiotemporal single-cell analysis we show that macrophage populations in mucosa and muscularis propria of the human colon contain multiple transcriptionally distinct subsets that display niche-specific localizations and functional properties. Our results also reveal tissue-specific cell-cell interactions and a limited number of TF that may be key players in the extensive monocyte-to-macrophage reprogramming and sub-tissular-specific localization observed.

Studies in mice show that macrophages in different organs display functional differences imprinted by tissue-specific cues from the local microenvironment (Guilliams et al., 2020; Okabe and Medzhitov, 2014). Moreover, it was recently shown that heterogeneous macrophages with different transcriptional programs are governed by sub-tissular niches (Chakarov et al., 2019). Therefore, to understand the functional diversity of macrophages within a tissue detailed spatiotemporal characterization of the macrophage population and their interactions with neighboring cells is needed. However, the nature of molecular signals that drive macrophage differentiation in the human gut is poorly characterized. Studies in humans and mice have clearly shown that mucosal macrophages in the intestine largely originate from bone marrow-derived monocytes (Bain et al., 2014; Bujko et al., 2018). Thus, probing colonic mucosal macrophages in histologically normal tissue is a unique possibility to study the monocyte-to-macrophage differentiation process in situ under steady state conditions.

Integration of our data suggests that circulating monocytes are elicited through post-capillary venules in the crypt area, after which some cells rapidly differentiate into different types of proinflammatory macrophages, whereas others migrate to the subepithelial region where they upregulate expression of genes related to endocytosis, wound healing and antigen presentation, while concomitantly downregulate proinflammatory genes. This latter subpopulation is thus ideally equipped to maintain mucosal barrier integrity with minimal collateral damage. Our cell-cell communication analysis suggested that the survival, migration and differentiation of LpM are controlled by an extensive interaction with tissue resident cells (endothelial cells, fibroblasts and epithelial cells) and immune cells. Stromal cells may provide a supply of macrophage-trophic factors (e.g. CSF-1, CSF3, IL-34), which are crucial for macrophage development and survival (Lavin et al., 2015), whereas several cell types expressed Notch ligands suggesting that the Notch-signaling pathway plays a role in monocyte-to-macrophage reprogramming (Sharma et al., 2020). Several receptor-ligand pairs have been shown to have inhibitory or anti-inflammatory effects on macrophages, in line with the idea that most tissue macrophages are tolerogenic under steady state conditions. Interestingly, however, many of these macrophage genes have been shown to promote tumor growth and may be targets for immunotherapy (e.g. SIRPA, SIGLEC10, AXL, MERTK, RIPK1) (Barkal et al., 2019; Li et al., 2021; Myers et al., 2019; Wang et al., 2018). Thus, strategies to target these genes to treat cancer should take into account possible side effects, such as drug-induced colitis, as observed by current checkpoint inhibition (Luoma et al., 2020).

Our gene regulatory network analysis identified cluster-specific TF that are likely to control subset-specific transcriptional programs, which strengthen the idea that the LpM population contains multiple transcriptionally stable macrophage subsets that coexist in the tissue. Interestingly, the transcriptomic profile of several LpM subsets is similar to macrophages associated with various inflammatory and fibrotic diseases (Martin et al., 2019; Qu et al., 2020; Ramachandran et al., 2019). This suggests that these disease-promoting macrophages also play a role under steady state conditions, but that the relative contribution of functionally different LpM subsets is important for maintenance of homeostasis.

Analysis of muscularis propria showed that many MM were positioned in close contact with nerves and cell-cell interaction analysis displayed numerous MM-neuron interactions. Such interactions included BMP2:BMP2R and CSF1R:IL34. As previously shown in mouse models (Gabanyi et al., 2016), this finding suggests that MM expressing BMP2 regulate peristaltic motility in the colon by activating BMP2R on enteric neurons, whereas neurons expressing IL34 feedback on CSF1R+ MM by stimulating their survival and differentiation. Most MM expressed high levels of C1Q genes and we identified interactions between genes of the complement system (C3:C3AR1, C5:C5AR2), suggesting that MM are involved in synapse pruning (Stephan et al., 2012). Finally, MM contained transcripts specific for Schwann cells indicating that MM phagocytose cellular material from such nerve-protecting cells. Together, these findings are in agreement with the concept that MM-neuron crosstalk play a pivotal role for gut homeostasis (De Schepper et al., 2018; Muller et al., 2014). MM were also positioned adjacent to vessels and the GO term positive regulation of angiogenesis was enriched in MM_1_, MM_5_ and MM_6_. Pseudotime trajectory analysis showed that these subsets were found along the “proinflammatory” branch, whereas neuron-associated MM subsets (first of all MM_11_) were linked to the “homeostatic” branch. This suggests that functionally specialized neuron- and blood vessel-associated MM coexist in the human colon to ensure proper functioning of enteric neurons and blood vessels, respectively (De Schepper et al., 2018).

Several studies in mice have suggested that macrophages important for vascular integrity selectively express Lyve-1 (Chakarov et al., 2019; Lim et al., 2018). However, we found that the vast majority of MM, both those associated with vessels and nerves, expressed Lyve-1 and Colec12, and we were not able to distinguish these subsets by in situ staining. We also identified Calprotectin+ MM scattered throughout the muscularis propria. Clustering and pseudotime trajectory analysis strongly suggested that the vast majority of TRM found in the muscularis propria originated from these incoming monocytes. Macrophages are the dominating leukocyte population in muscularis propria and submucosa. Interestingly to this end, we found that MM subsets and SmM expressed high levels of several monocyte-attracting chemokines (e.g. CCL3, CCL4, CCL3L1, CCL4L2), suggesting that TRM in these compartments are important for continuous recruitment of monocytes.

Somewhat surprisingly, we found that subpopulations of LpM and MM showed phenotypic similarities with embryonic-derived macrophages in mouse colon (De Schepper et al., 2018) and with colonic macrophages in human fetuses (Fawkner-Corbett et al., 2021). As far as we know there are no established markers to separate embryonic-derived and from bone marrow monocyte-derived macrophages in humans, and evidence to suggest that human macrophages in adult life originate from embryonic precursors is sparse. We and others have shown that macrophages in various tissues may live for several years (Eguiluz-Gracia et al., 2016; Patel et al., 2021). However, the origin of these long-lived cells has not been determined. Although the fraction of macrophages with an “embryonic” signature in our study was minor, it is possible that embryonic-derived macrophages play a more prominent role other settings, in particular during infancy. Thus, it should be studied further whether genes such as DNASE1L3 and ADAMDEC1 could be useful markers to distinguish macrophage lineages.

Together, we find that LpM and MM are extremely heterogeneous and consist of several subsets with distinct functional properties. It appears that maintenance of homeostasis depends on coexistence of both proinflammatory and protective/homeostatic subtypes. Our data also give insights into cell-cell interactions and key TF that are likely to control tissue-specific macrophage reprogramming. This work constitutes an important framework to understand the complexity of macrophage biology in the human gut and to identify potential targets to better treat inflammatory disorders and cancer in the future.

## Acknowledgments

We thank all patients and staff at Akershus University Hospital for providing tissue samples. Dr Susanne Lorenz at the Genomics Core Facility at Oslo University Hospital is greatly acknowledged for supervising the sequencing. We also thank Dr Frode Skjeldal at the Oslo NorMIC imaging platform at the Department of Biosciences, University of Oslo, for expert help with confocal microscopy analysis. Kjersti Thorvaldsen Hagen is greatly acknowledged for technical help with tissue processing and immunostainings. This work was supported by the Norwegian Cancer Society and Southern and Eastern Norway Regional Health Authority.

## STAR METHODS

### KEY RESOURCES TABLE

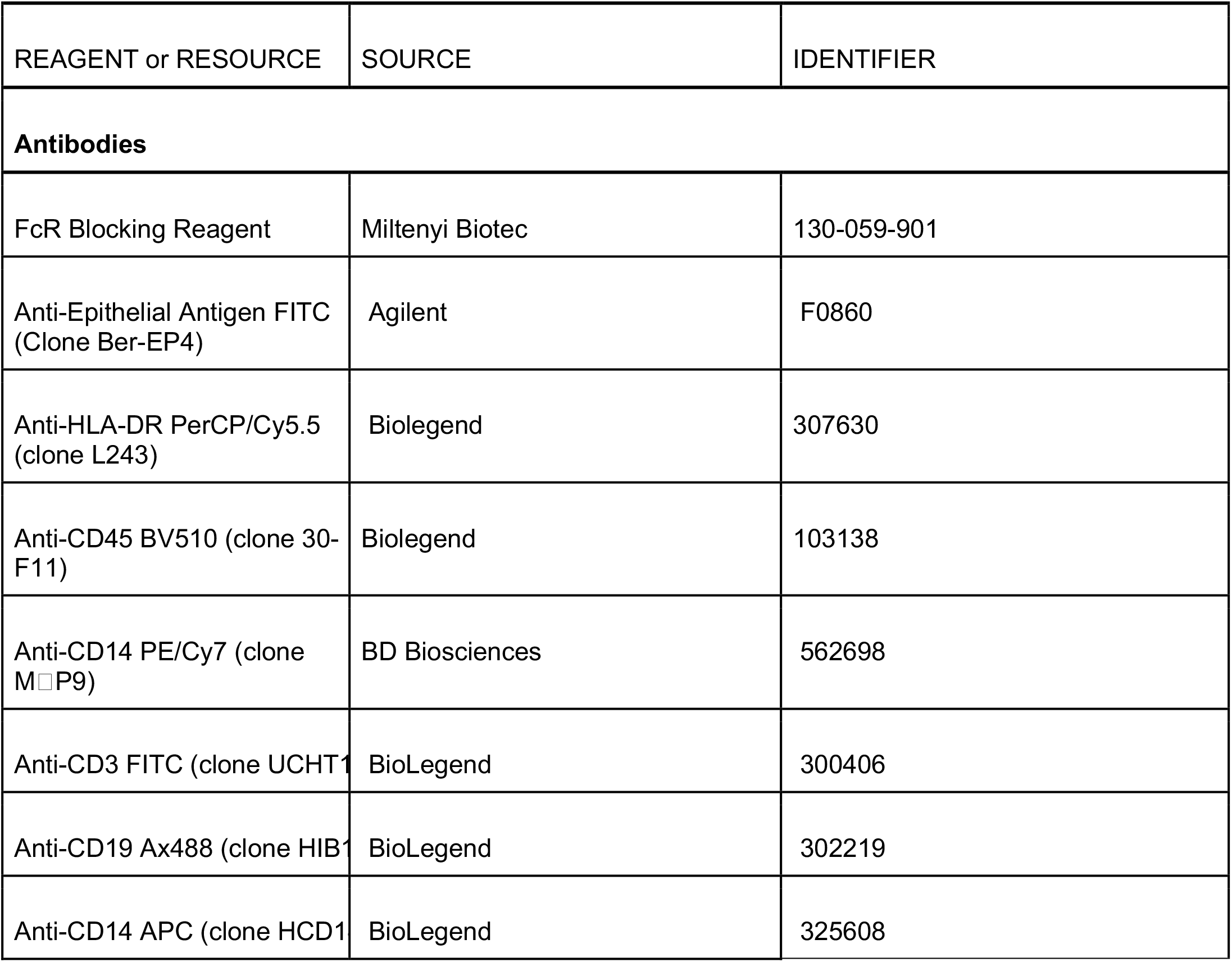
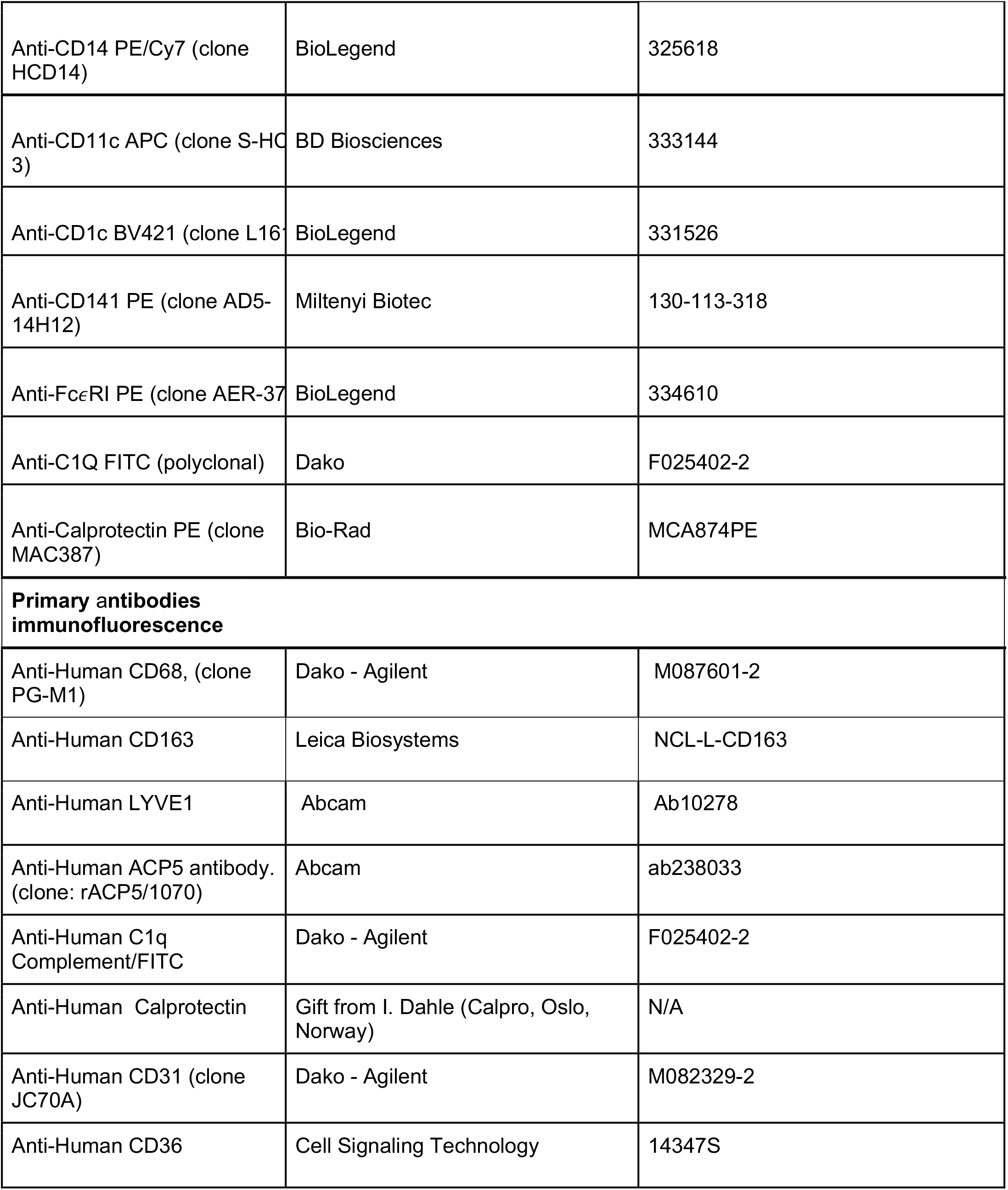
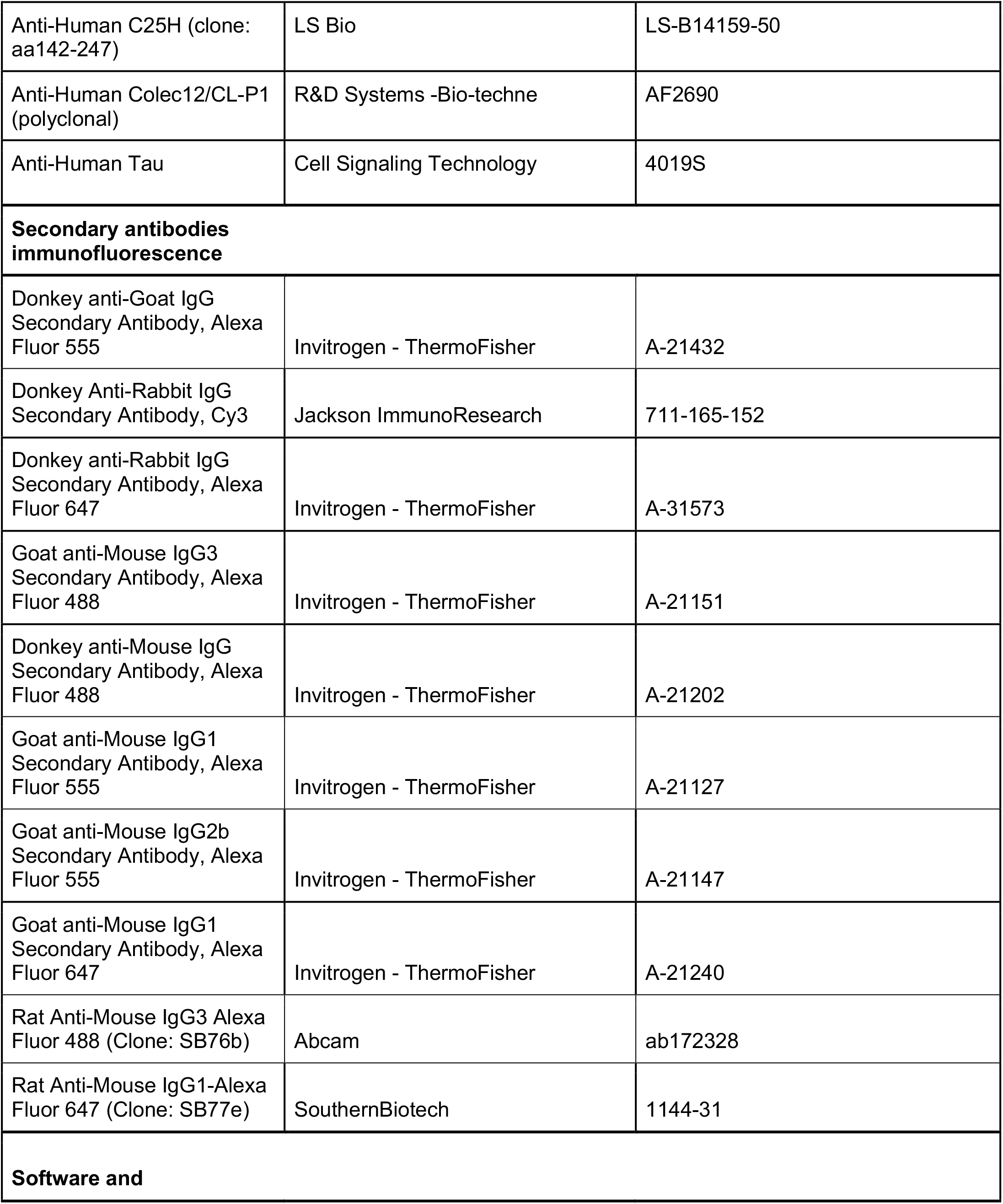
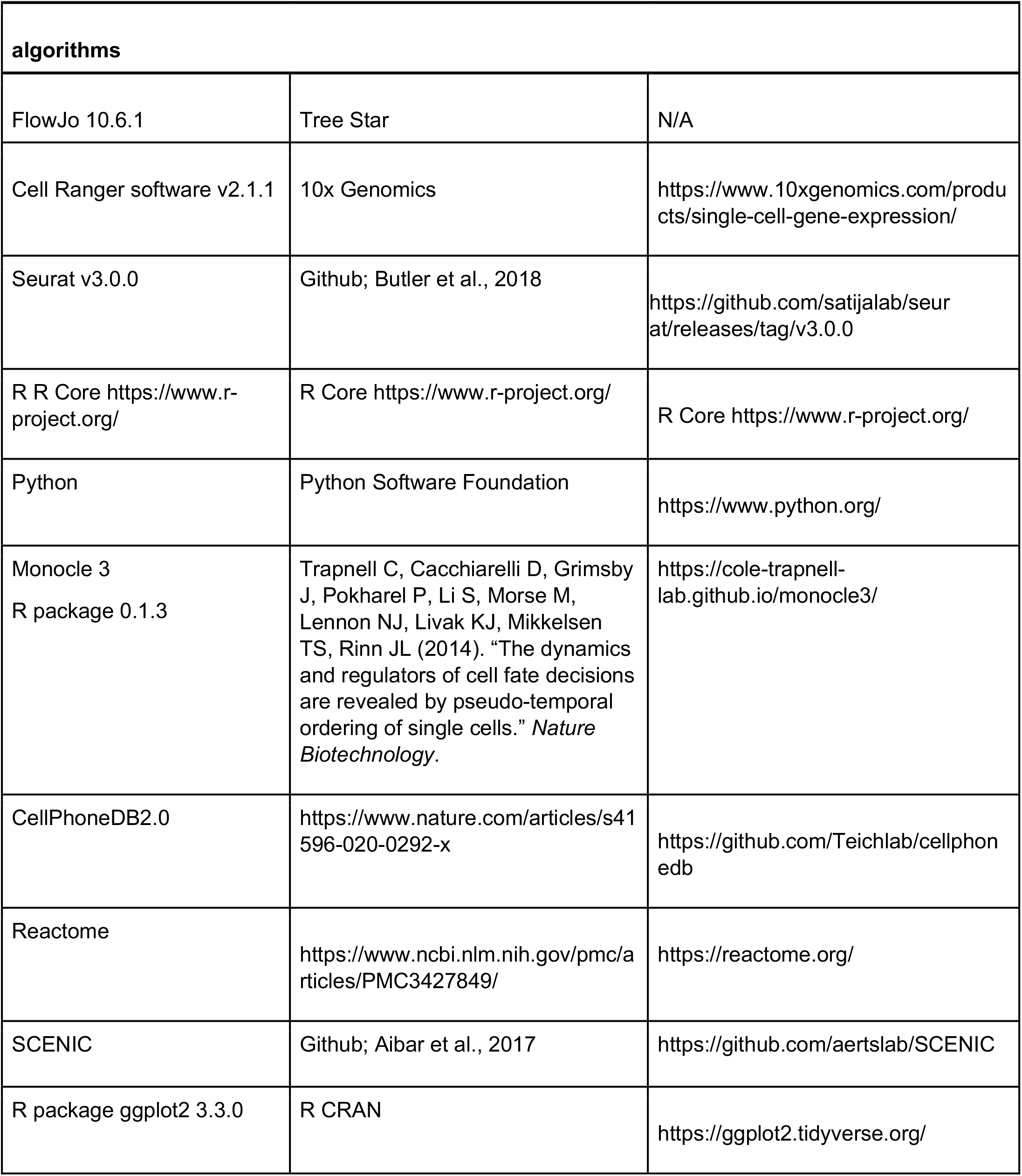
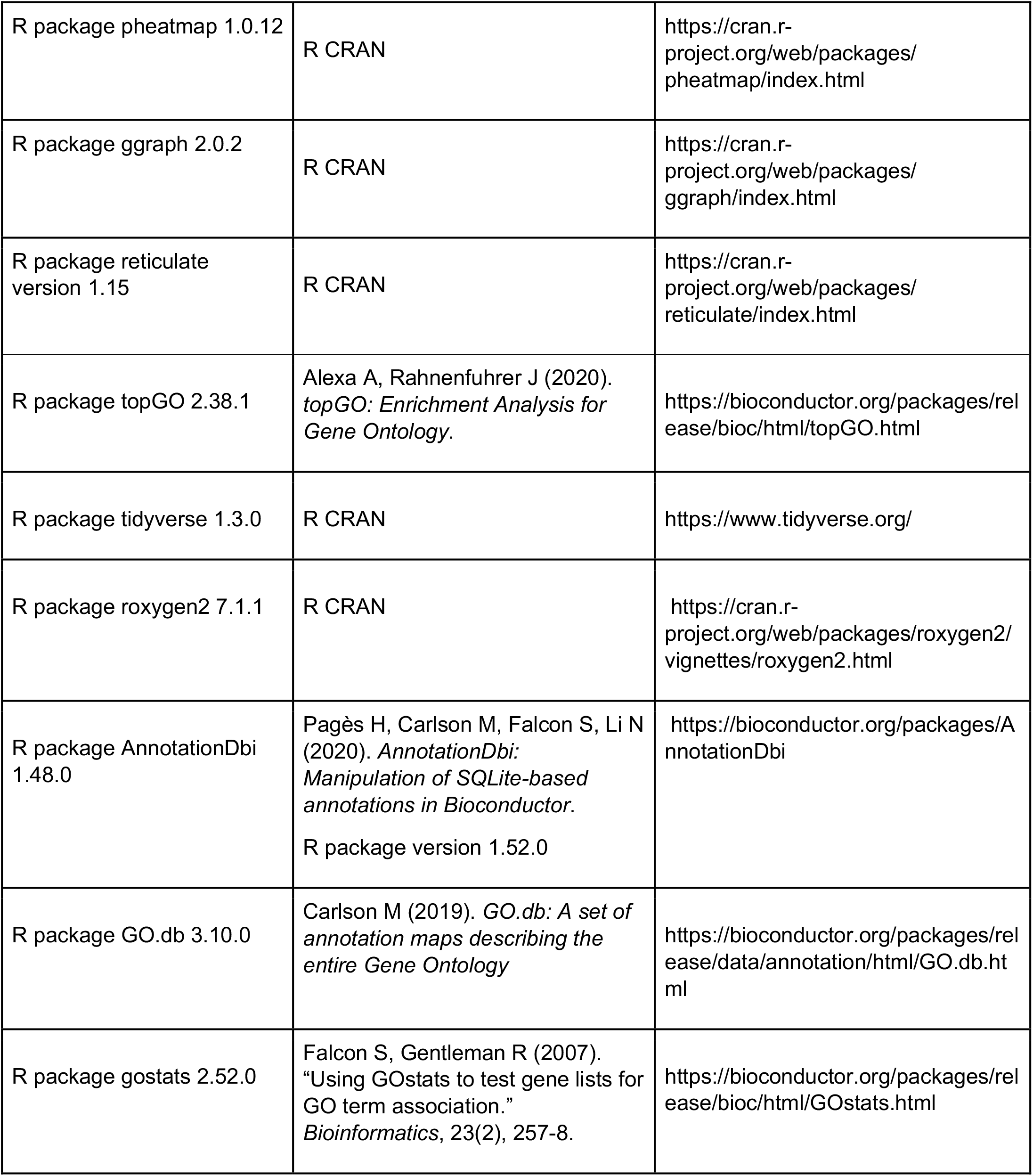
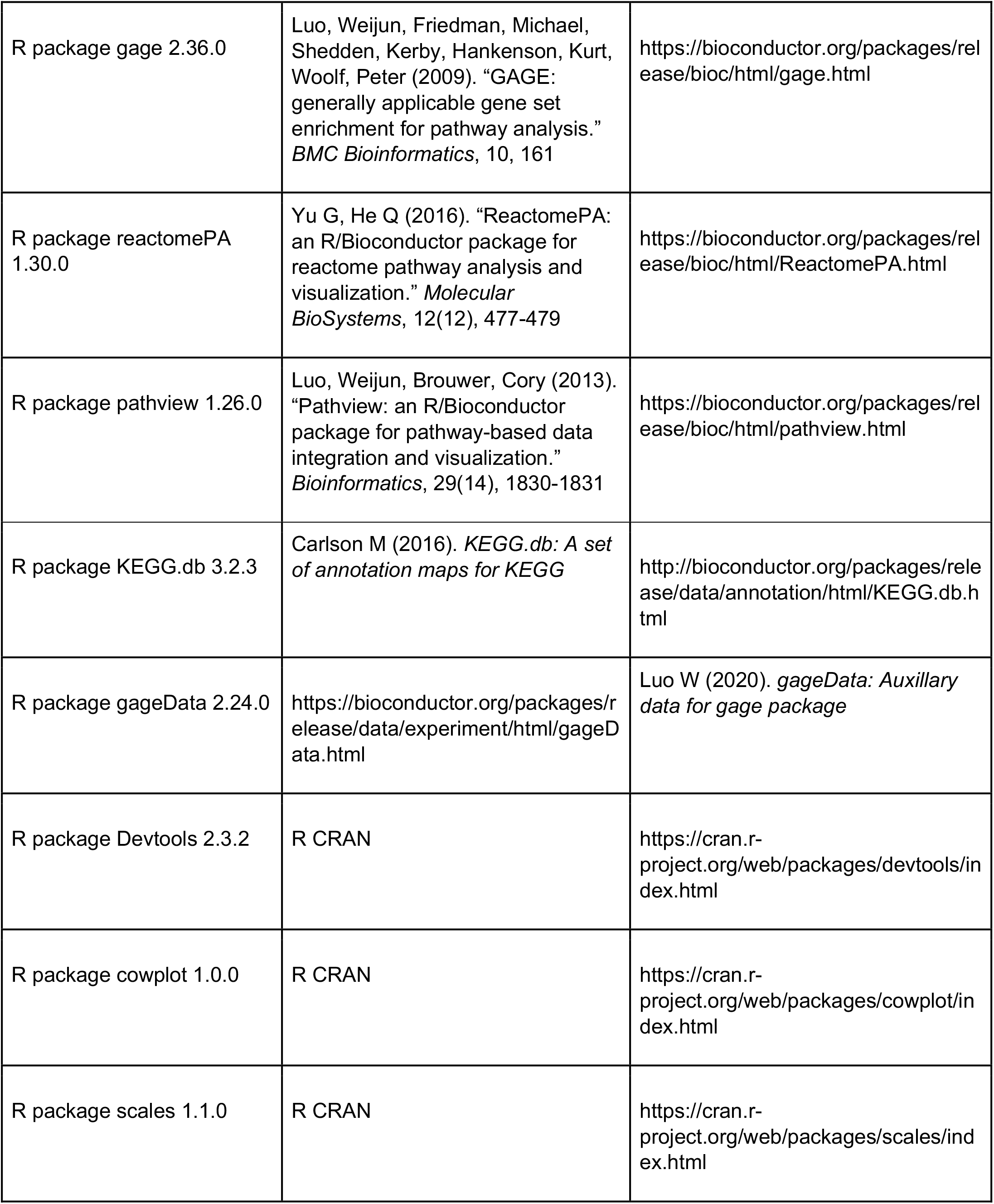
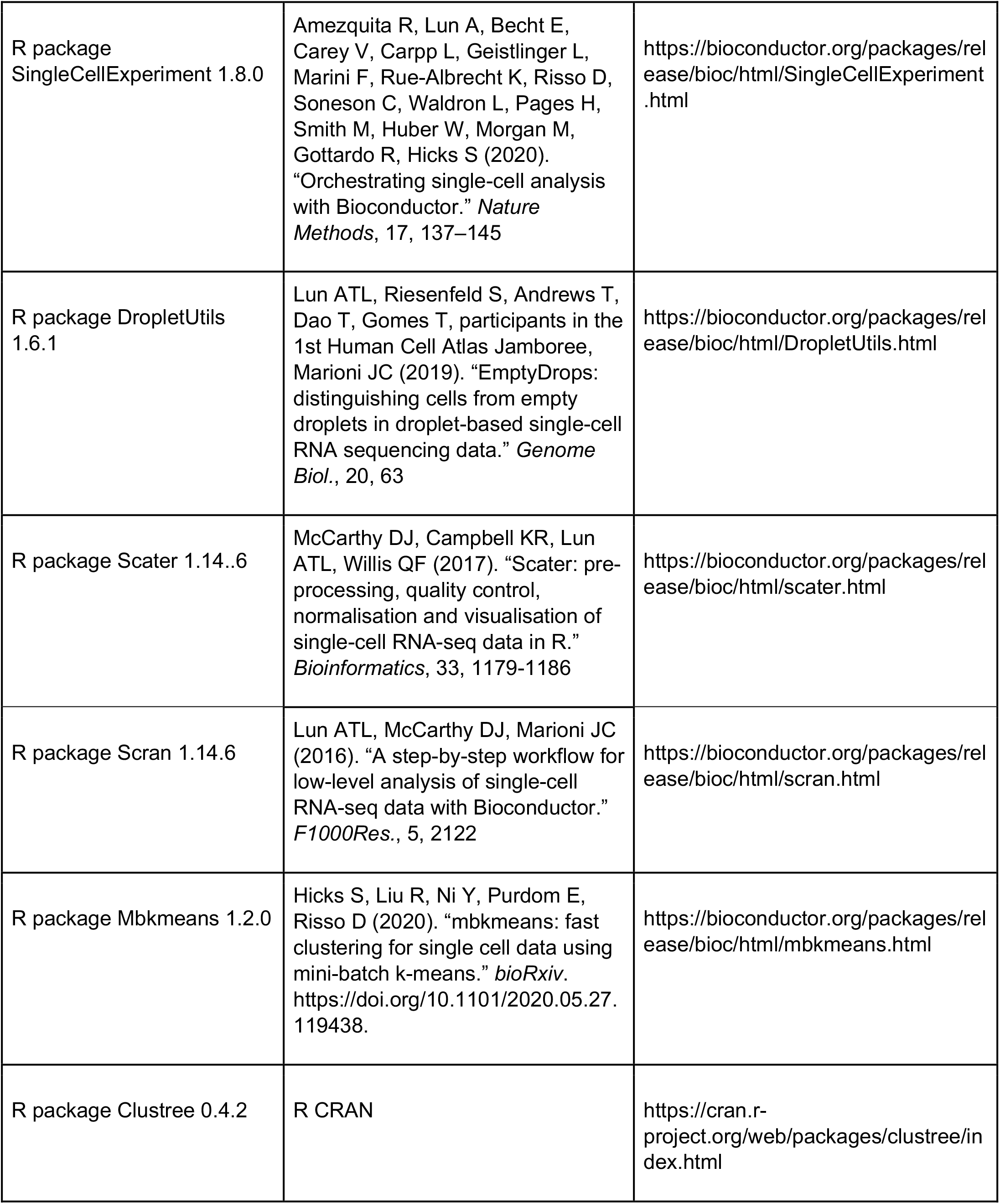
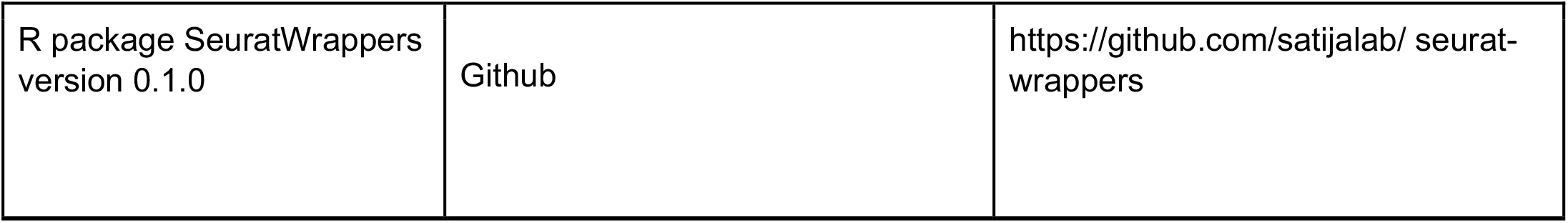

### Materials Availability

This work did not generate new unique reagents.

### Data and Code Availability

Single cell RNA-Seq datasets generated in this study are deposited in the Genome Expression Omnibus under the following accession numbers: to be updated.

Reference datasets used in this study:

Human UC scRNA-seq dataset (Smillie et al., 2019) SCP:SCP259

Human ENS scRNAseq dataset (Drokhlyansky et al., 2020) SCP: SCP1038

Human fetal intestinal scRNA-seq dataset (Fawkner-Corbett et al., 2021) GEO:GSE158702

## EXPERIMENTAL MODEL AND SUBJECT DETAILS

### Patients and tissue samples

Colonic resections were obtained from patients operated for sigmoid colon cancer at Akershus University Hospital (Ahus, 1478 Lørenskog, Norway). The resected colon was immediately examined by an experienced pathologist and macroscopically normal colon, at least 10 cm from the tumor, were placed in vials with RPMI 1640 and put on ice for transport. The study was performed in accordance with the Declaration of Helsinki. Written informed consent was obtained from all participants, and the study was approved by the Regional Committee for Medical Research Ethics (REK, 2018/703, Health Region South-East, Norway). For scRNA-seq, 4 colon specimens from sigmoid or ascending colon were obtained (age 62-78, 3 males). Immunofluorescence stainings were done on samples from 8 patients (age 62-78, 4 males) from ascending, transverse, descending and sigmoid colon. None of the patients had autoimmune, infectious or inflammatory diseases, nor received neoadjuvant chemotherapy or radiation therapy before the operation. All patients were operated following the national guidelines.

## METHOD DETAILS

### Single cell dissociation

Resected colonic tissues were processed within 2 h after removal from the patient. Single cell suspensions of colonic resections were obtained using a modified version of a previously published protocol (Bujko et al., 2018). The intestinal specimens were opened longitudinally and washed in Dulbecco’s phosphate-buffered saline (PBS). The muscularis propria was first removed with scissors, after which the mucosa was dissected in narrow strips. The mucosal fragments were then incubated with shaking in PBS with 2 mM EDTA (Sigma-Aldrich) and 1% FCS (Sigma-Aldrich) three times for 15 min at 37°C. The remaining tissue was minced and digested with stirring for 60 min in complete RPMI (RPMI1640; Lonza; supplemented with 10% fetal calf serum (FCS, 1% Pen/Strep; Lonza) containing 0.25 mg/ml Liberase TL (Roche) and 20 U/ml DNase I (Sigma). Digested cell suspension was passed through a 100-μm filter and washed.

### Flow cytometry and cell sorting

Released tissue cells were stained in aliquots of 1 ×10^6^ cells/100 μL of PBS with 2% fetal calf serum (FCS, GIBCO) and 0.1% NaN3 for 30min on ice. Non-specific staining was blocked with 10 μL FcR Blocking Reagent (Miltenyi Biotec) prior to staining. Dead cells were excluded by TO-PRO™-1 Iodide (ThermoFisher) or Fixable Viability Dye eFluor 780 (eBioscience) staining. Analysis was performed with a BD LSRFortessa X20 and sorting with a FACS Aria IIIu (BD Biosciences) running BD FACSDIVA 9.0 software. Purity of > 98% was achieved in sorted populations. Data were processed with FlowJo 10.6.1 (Tree Star, Inc). Intracellular staining was performed after surface staining, using a Fixation & Permeabilization Buffer Set (eBioscience) according to manufacturer’s instructions. A full list of antibodies is provided in the Key Resources Table.

### Single Cell RNA-sequencing RNaseq libraries preparation

Cellular suspensions (~15000 cells, with expected recovery of ~7500 cells) of sorted CD45+ HLA-DR+ CD14+ macrophages from colonic mucosa and muscularis propria were loaded on the 10X Chromium Controller instrument (10X Genomics) according to the manufacturer’s protocol using the 10X GEMCode proprietary technology. All samples from individual patients were loaded in one batch. The Chromium Single Cell 3’ v2 Reagent kit (10X Genomics) was used to generate the cDNA and prepare the libraries, according to the manufacturer’s protocol. The libraries were then equimolarly pooled and sequenced on an Illumina NextSeq500 using HighOutput flow cells v2.5. A coverage of 400M reads per sample was targeted, in order to obtain 50 000 reads per cell. The raw data were then demultiplexed and processed with the Cell Ranger software (10X Genomics) v2.1.1.

### Pre-processing scRNA-seq data

In total, we analyzed 63917 human cells from donors (n = 4). We aligned the reads of the input dataset to the GRCh38 reference genomes, and estimated cell-containing partitions and associated unique molecular identifiers (UMIs) using the Cell Ranger Chromium Single Cell RNA-seq version 3.0.2. We performed data preparation using Seurat R packages. Genes expressed in fewer than 3 cells in a sample were excluded, as well as cells that expressed fewer than 200 genes and mitochondrial gene content > 5% of the total UMI count. We normalized data by using gene counts for each cell that were divided by the total counts for that cell and multiplied by 10000 and then log-transforming. Subsequently, we identify genes that are outliers on a ‘mean variability plot’ using the vst method with 2000 genes. For mucosa and muscularis data, we separately found integration anchors and then performed data integration using a pre-computed anchorset with default parameters. Finally, we scaled data and centered genes in the dataset using linear model.

### Dimensionality reduction, clustering and differential expression analysis

We ran PCA dimensionality reduction with 30 PCs to compute and store (on 2000 variable genes). We estimated dimensions of reduction parameter (for LpM equal 13 and for MM type equal 20) and constructed a Shared Nearest Neighbor (SNN) Graph for given datasets. We first determined the k-nearest neighbors of each cell. We used this k-Nearest Neighbour graph to construct the SNN graph by calculating the neighborhood overlap (Jaccard index) between every cell and its 20 nearest neighbors. To obtain the resolution parameter we used clustree from the clustree R package with resolution varying from 0.1 to 2.0. We got resolution parameter for LpM equal 0.7 and for MM 0.6.

We then ran the Uniform Manifold Approximation and Projection (UMAP) dimensional reduction technique with PCA dimension reduction and we found differentially expressed genes for each of the clusters in the datasets.

We identified non-macrophage cell types such as B-cells and T-cells and these were filtered out. We got 5263 cells from mucosa and 14649 cells for MM. Then we ran UMAP and we found differentially expressed genes for each of the clusters in the datasets.

All heat maps, UMAP visualizations, violin plots and dot plots were produced using Seurat functions in conjunction with the ggplot2, pheatmap and grid R packages. ClusterMap (Gao et al., 2019) was used to compute a similarity metric for subclusters between LpM and MM.

### Developmental trajectory inference and transcriptional regulation

To generate pseudotemporal dynamics we used the Monocle R package. We ordered cells in a semi-supervised manner on the basis of their Seurat clustering, scaled the resulting pseudotime values from 0 to 1, and mapped them onto UMAP visualizations generated by Seurat. Differentially expressed genes along this trajectory were identified using the Moran’s I test. For transcription factor analysis, we obtained a list of all genes identified as human transcription factors. To analyze transcription factor regulons further, we adopted the Single Cell Regulatory Network Inference and Clustering, (SCENIC) (https://aertslab.org/#scenic), using default parameters and the normalized data matrices from Seurat as input. SCENIC is a combination of 3 packages (GENIE3, RcisTarget and AUCell). For motif visualization we obtained the highest normalized enrichment score of the motif in the gene-set.

### Identification of significant ligand-receptor pairs

For comprehensive systematic analysis of inter-lineage interactions, we used CellPhoneDB2.0 (https://www.cellphonedb.org). CellPhoneDB2.0 is a manually curated repository of ligands, receptors and their interactions, integrated with a statistical package for inferring cell–cell communication networks from single-cell transcriptomic data. This package searches for ligand-receptor interactions and outputs multiple result files based on curated databases such as UniProt, IUPHAR, and Ensembl.

Each dataset was analyzed using matrices from Seurat and datasets covering epithelial, endothelial, fibroblast, immune cells (Smillie et al., 2019) subsets and enteric neurons (Drokhlyansky et al., 2020). Significant ligand-receptor pairs identified from datasets, with adjusted p value < 0.05 were extracted, requiring the ligand and receptor to be expressed in at least 10% of the cells.

### Immunofluorescence stainings

Sections of formalin-fixed and paraffin-embedded tissue were cut in series at 4 μm, mounted on Superfrost Plus object glasses (Thermo Fisher Scientific), and washed sequentially in xylene, ethanol, and PBS. Heat-induced epitope retrieval was performed by boiling sections for 20 min in citrate buffer (pH 6.0) and cooled to room temperature before staining. Sections were incubated with mixtures of primary antibodies for 1 h at 37°C, rinsed in PBS, and incubated with secondary antibodies for 1.5h at RT. Sections were then incubated for 5 min at RT in Hoechst 33342 nucleic acid stain, and stained sections were mounted with ProLong Glass Antifade mountant (Molecular Probes). Laser scanning confocal microscopy was performed by acquiring tile scans on an Andor Dragonfly equipped with a fusion stitcher. The Andor Dragonfly was built on a Nikon TiE inverted microscope equipped a 60x/1.40 NA oil immersion objective.

---

**Figure S1:**
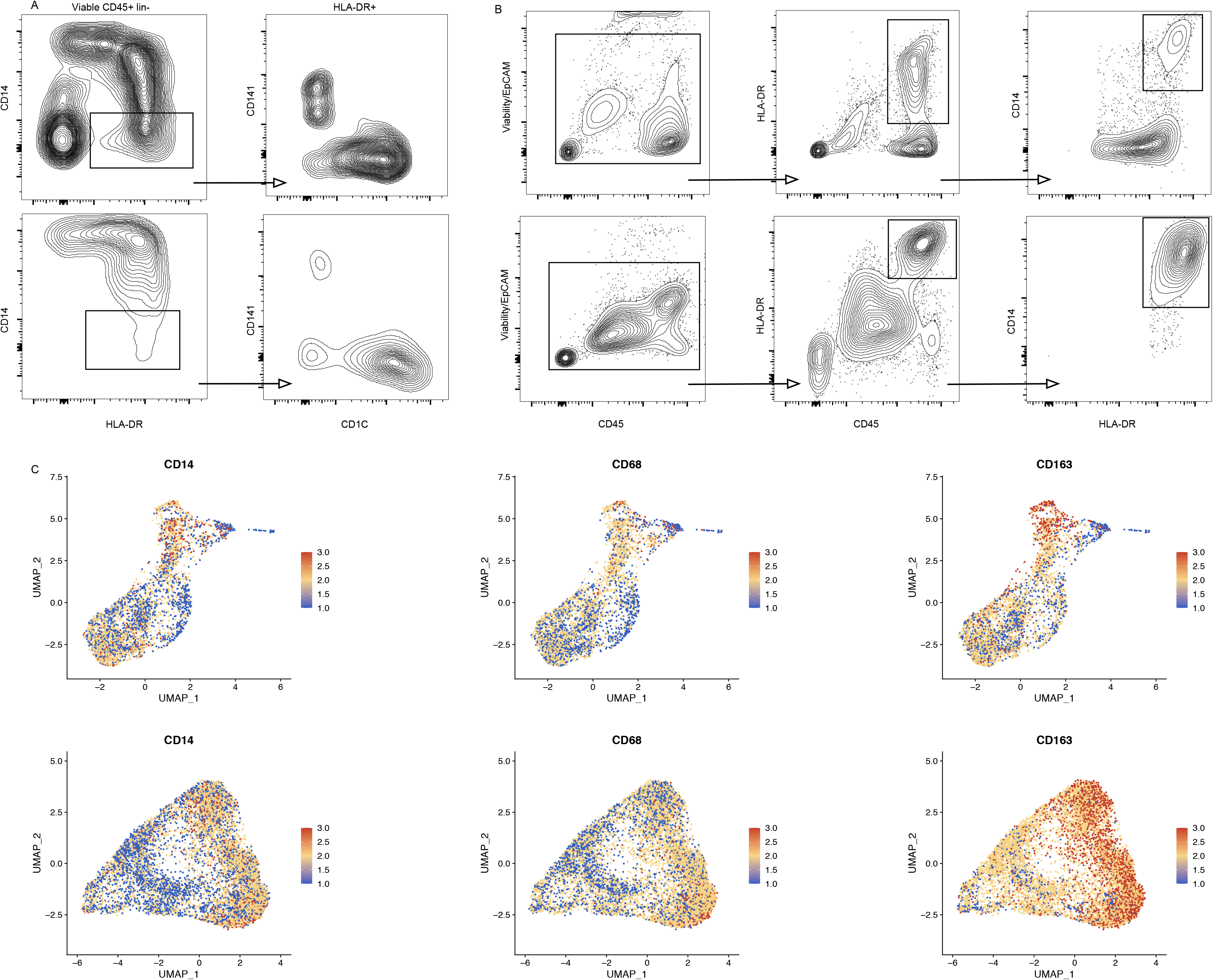
Characterization and flow cytometry sorting strategy for LpM and MM. Representative flow cytometry plots to identify CD14+ macrophages and cDC in colon LP (top) and muscularis propria (bottom) (A) and sorting strategy (B) for LpM (top) and MM (bottom). (C) UMAP plots of selected macrophage-related genes in LpM (top) and MM (bottom).

**Figure S2:**
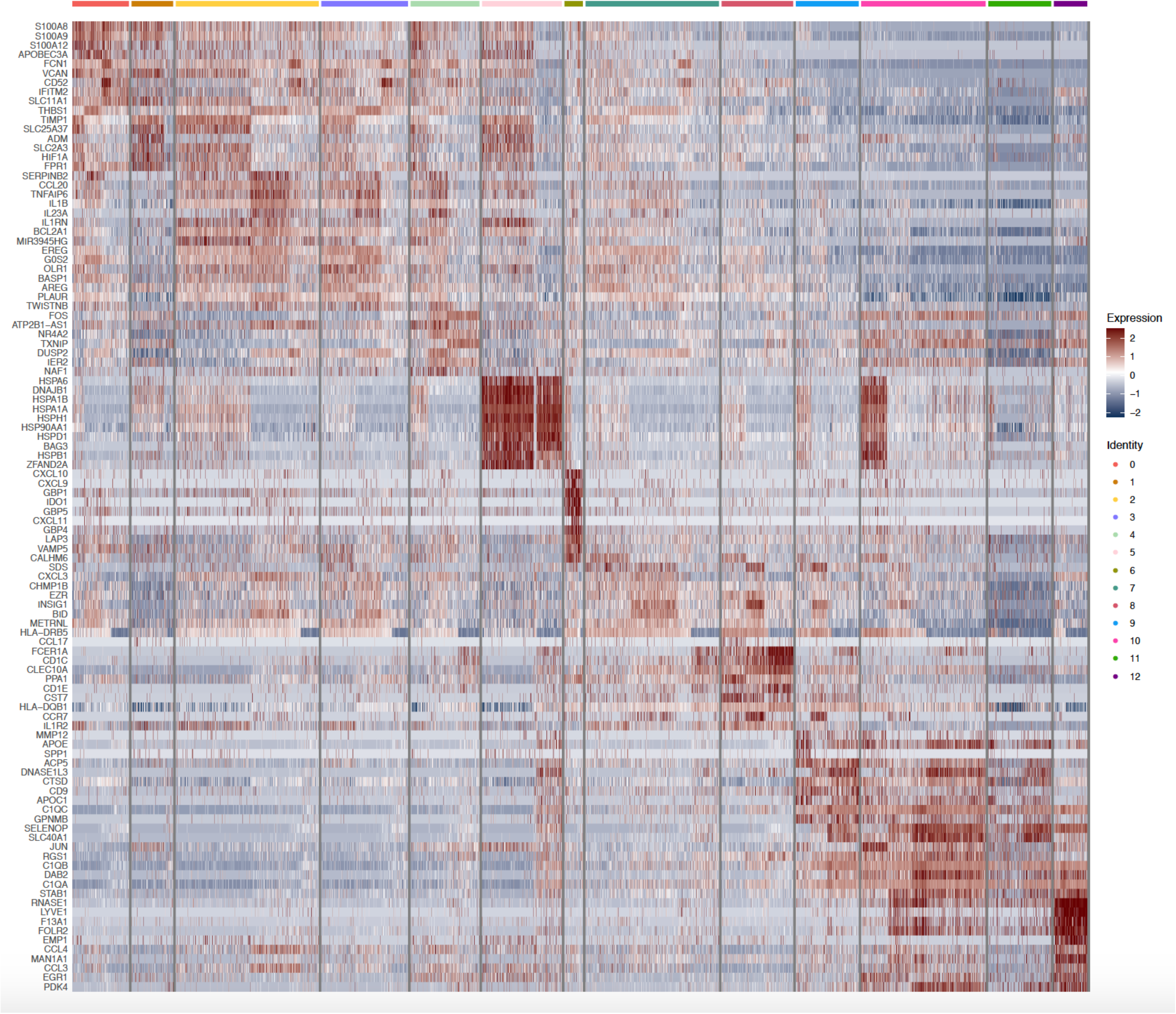
Heatmap of top 10 differentially expressed genes in clustered LpM.

**Figure S3:**
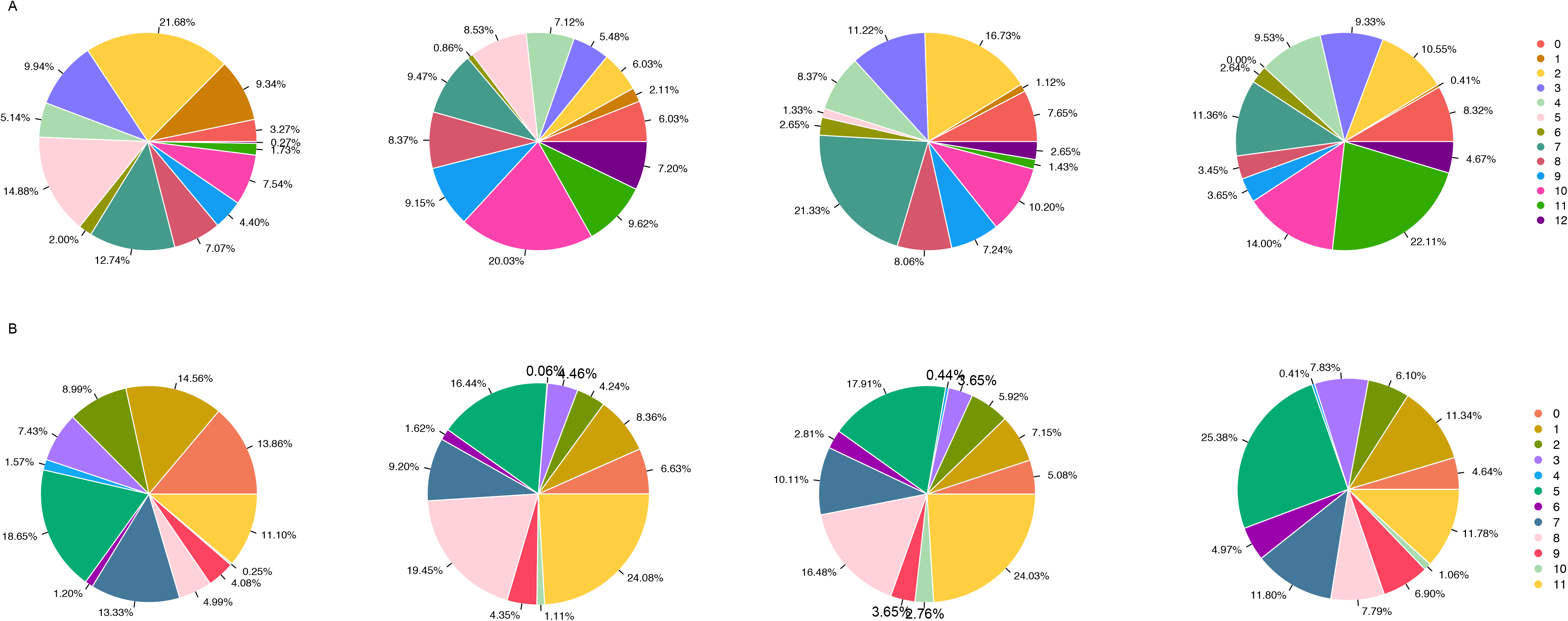
Representation of cell frequency from included donors in the clusters. A) LpM and B) MM.

**Figure S4.**
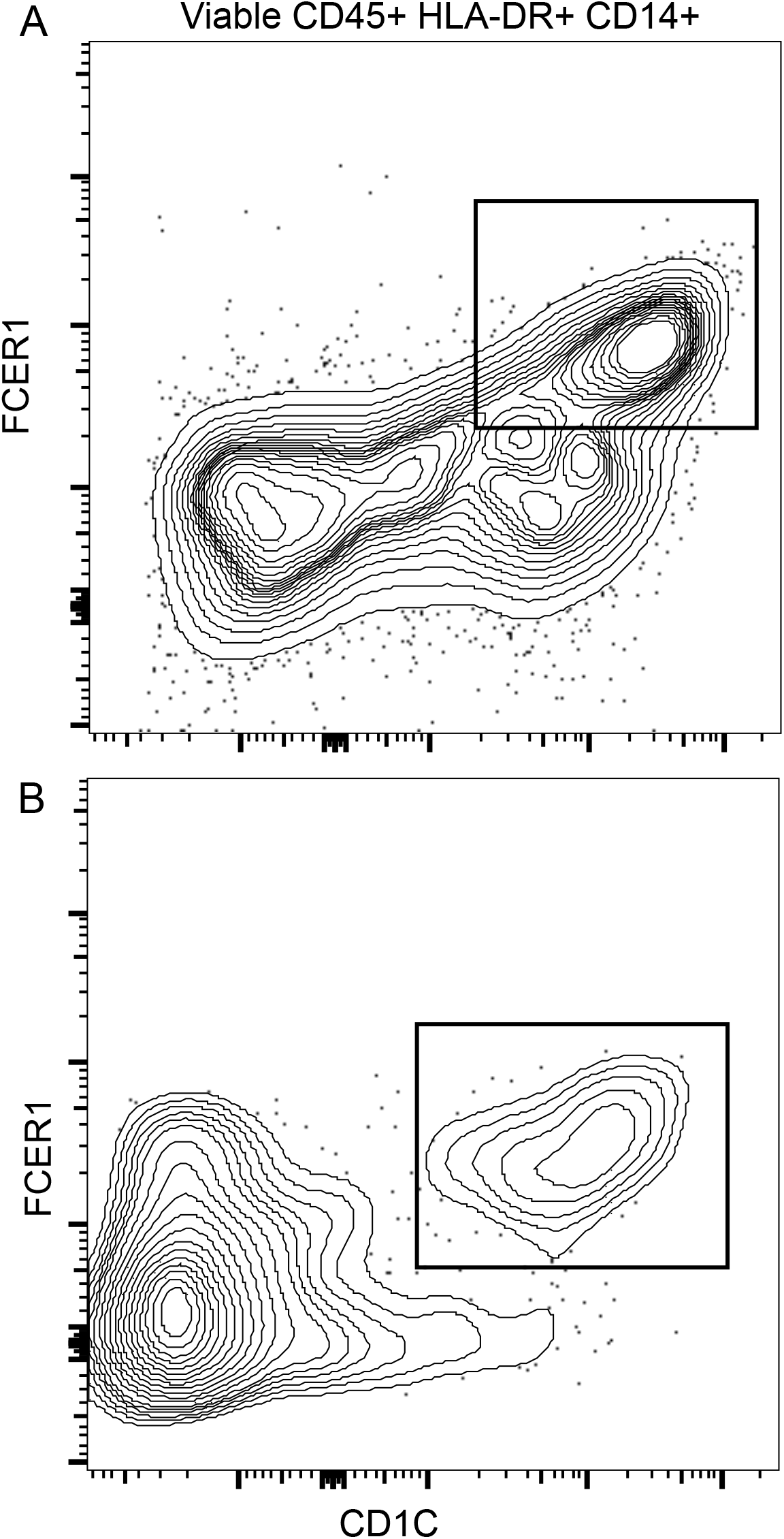
Identification of DC3-like cells in colon. Representative flow cytometry plots showing CD14^+^CD1c^+^FcER1^+^ cells in A) mucosa and B) muscularis propria.

**Figure S5:**
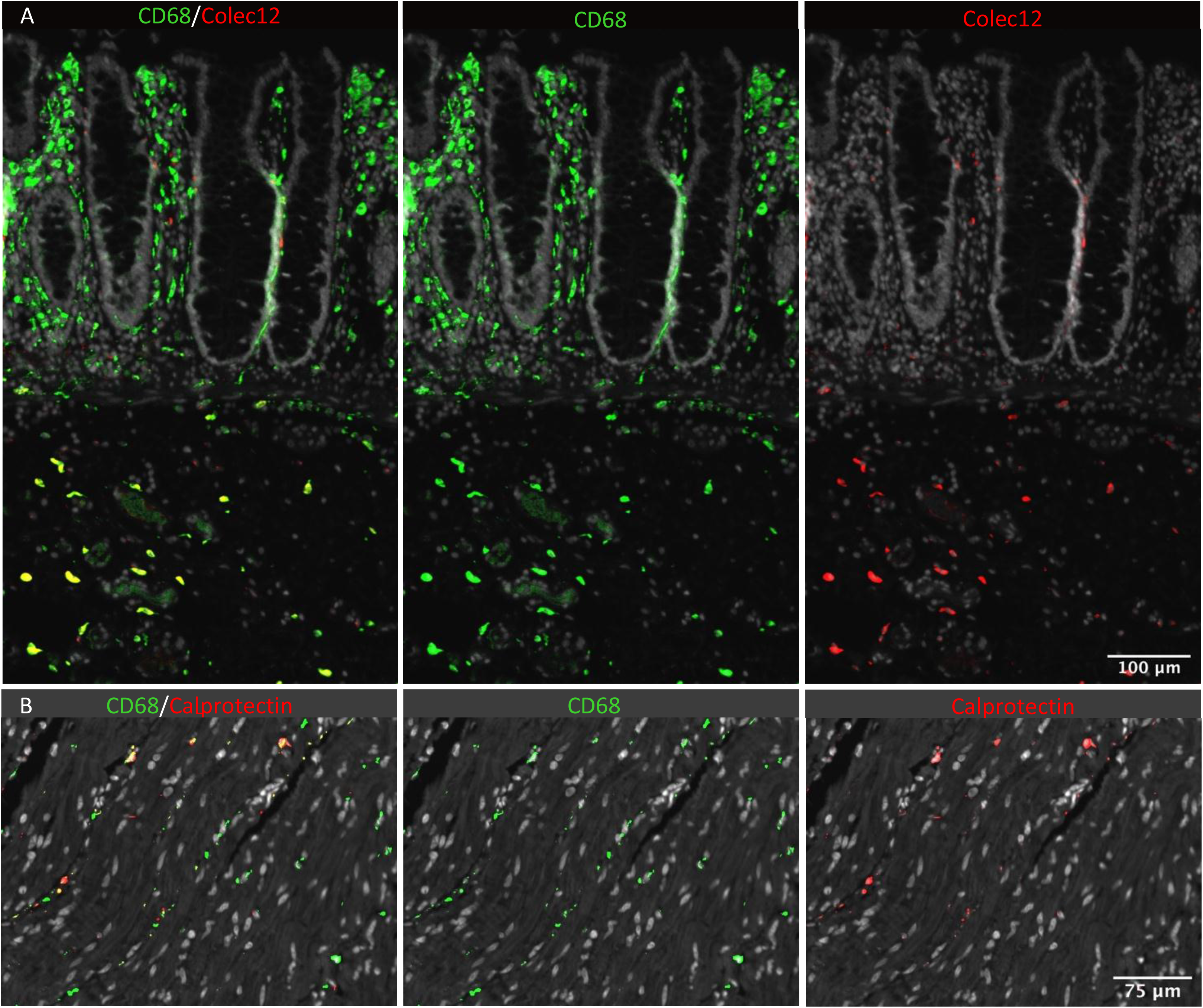
In situ localization of macrophages in human colon. A) Section of mucosa and submucosa stained for CD68 (green) and Colec12 (red), B) section of muscularis stained for CD68 (green) and calprotectin (S100A8/S100A9) (red). Sections were counterstained with Hoechst DNA-stain (gray). Representative of n≥3

**Figure S6:**
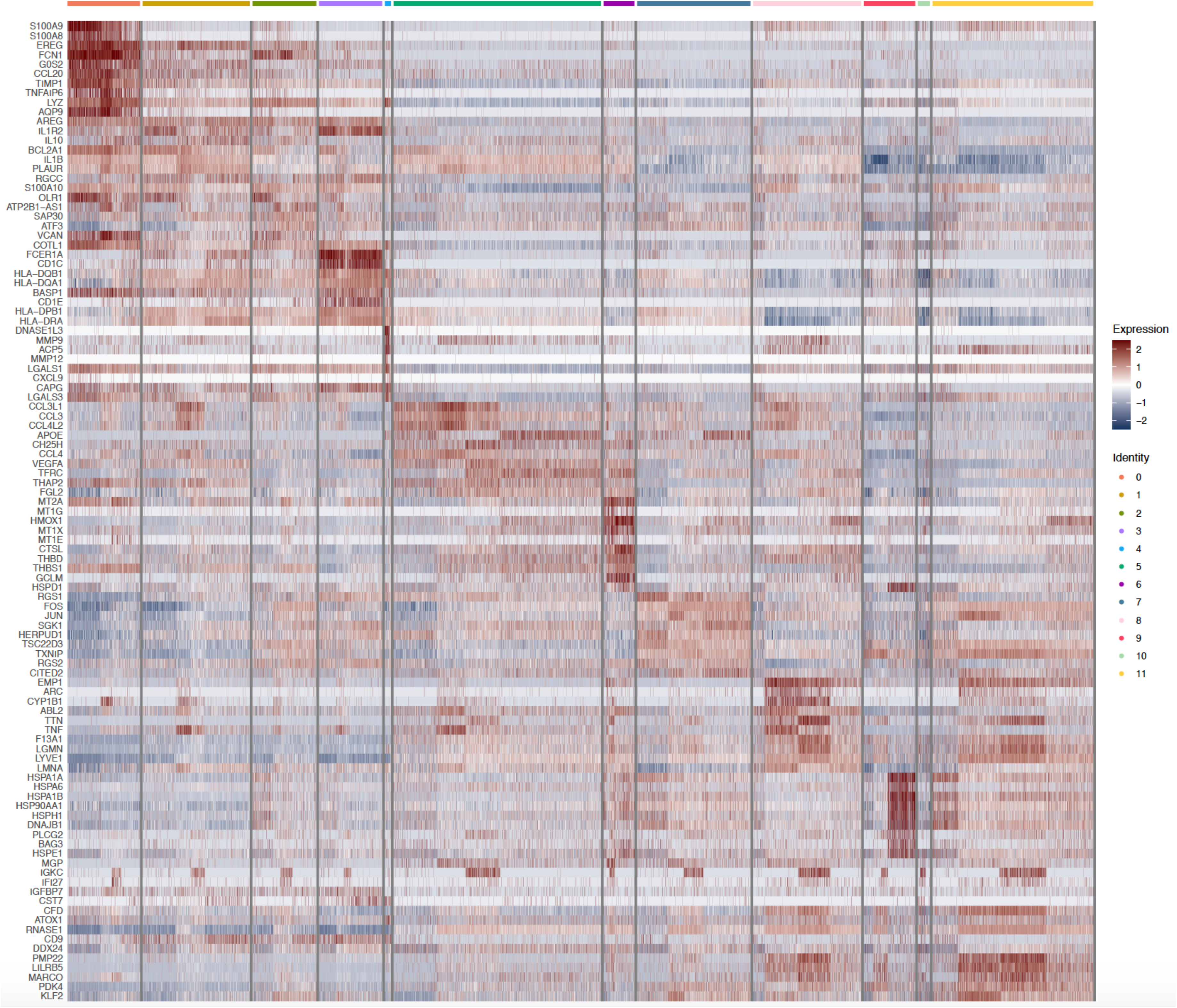
Heatmap of top 10 differentially expressed genes in clustered MM.

**Figure S7:**
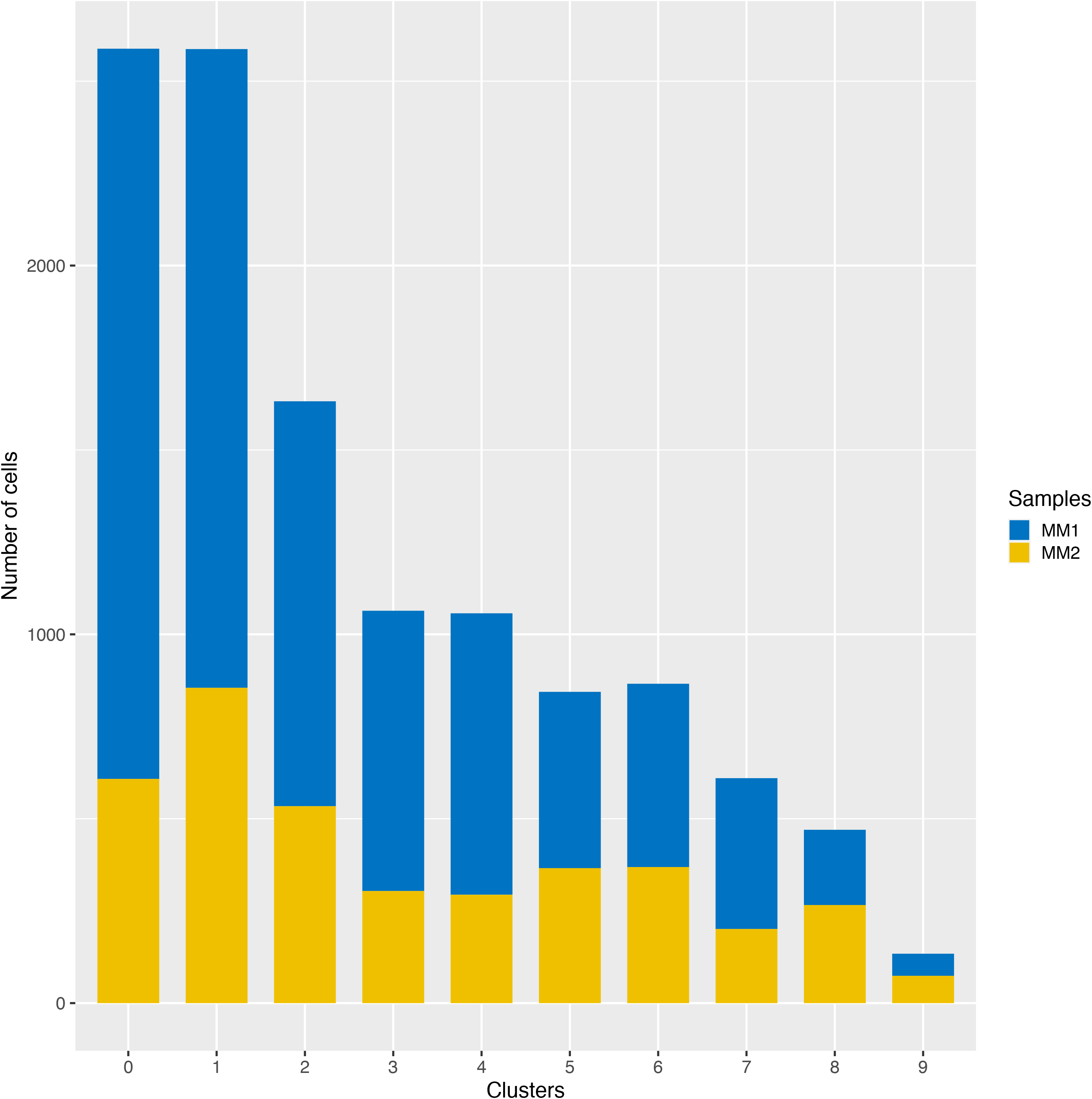
Representation of cells in MM clusters obtained from two sites (MM1, MM2) of muscularis propria (n=3).

**Figure S8:**
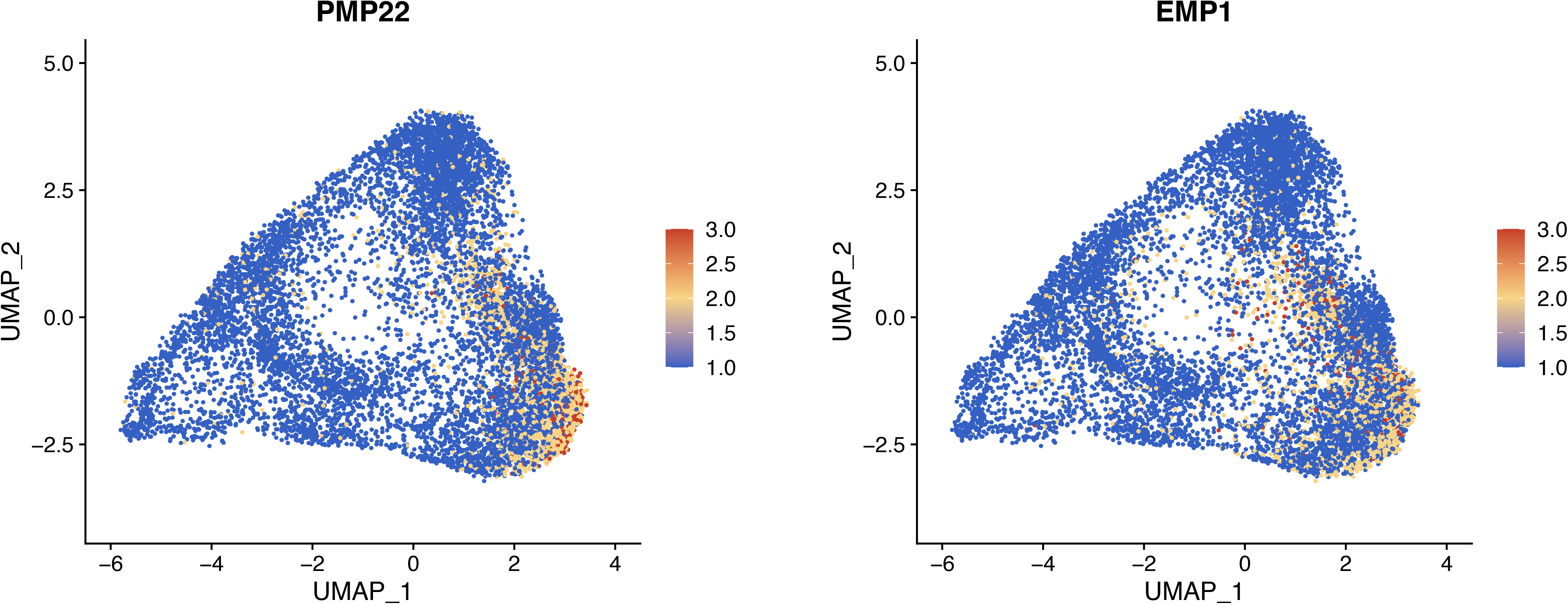
UMAP plot of MM showing the expression of PMP22 and EMP1.

**Figure S9:**
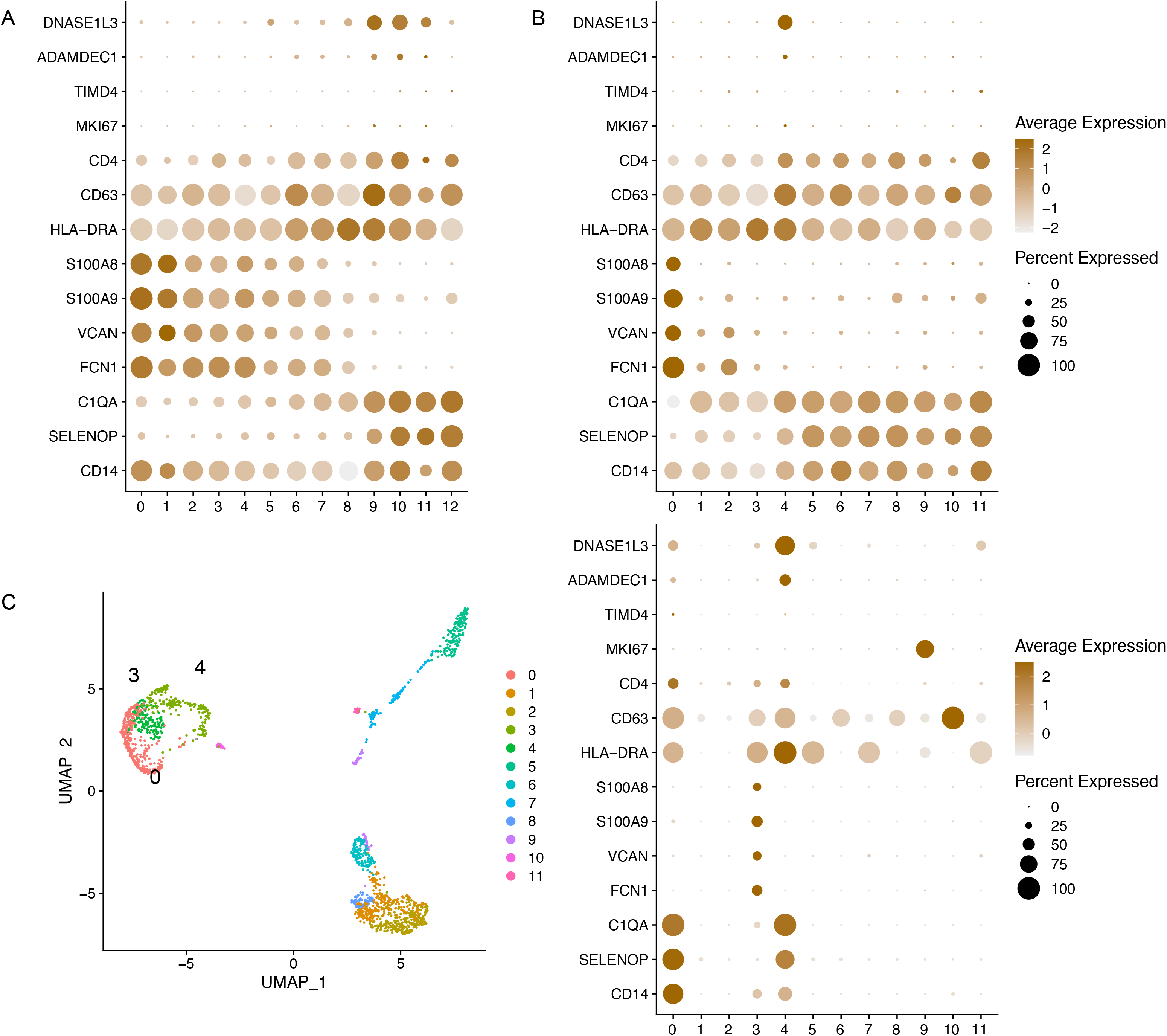
Expression of genes associated with embryonic-derived macrophages. Dot plot of average expression of selected genes in LpM (A) and MM (B). C) UMAP plot of intestinal-derived immune cells (left, macrophage clusters indicated (0, 3, 4)) and dot plot of average expression in fetal macrophage clusters (right). Intestinal samples are from post conception week 12-22.

**Figure S10:**
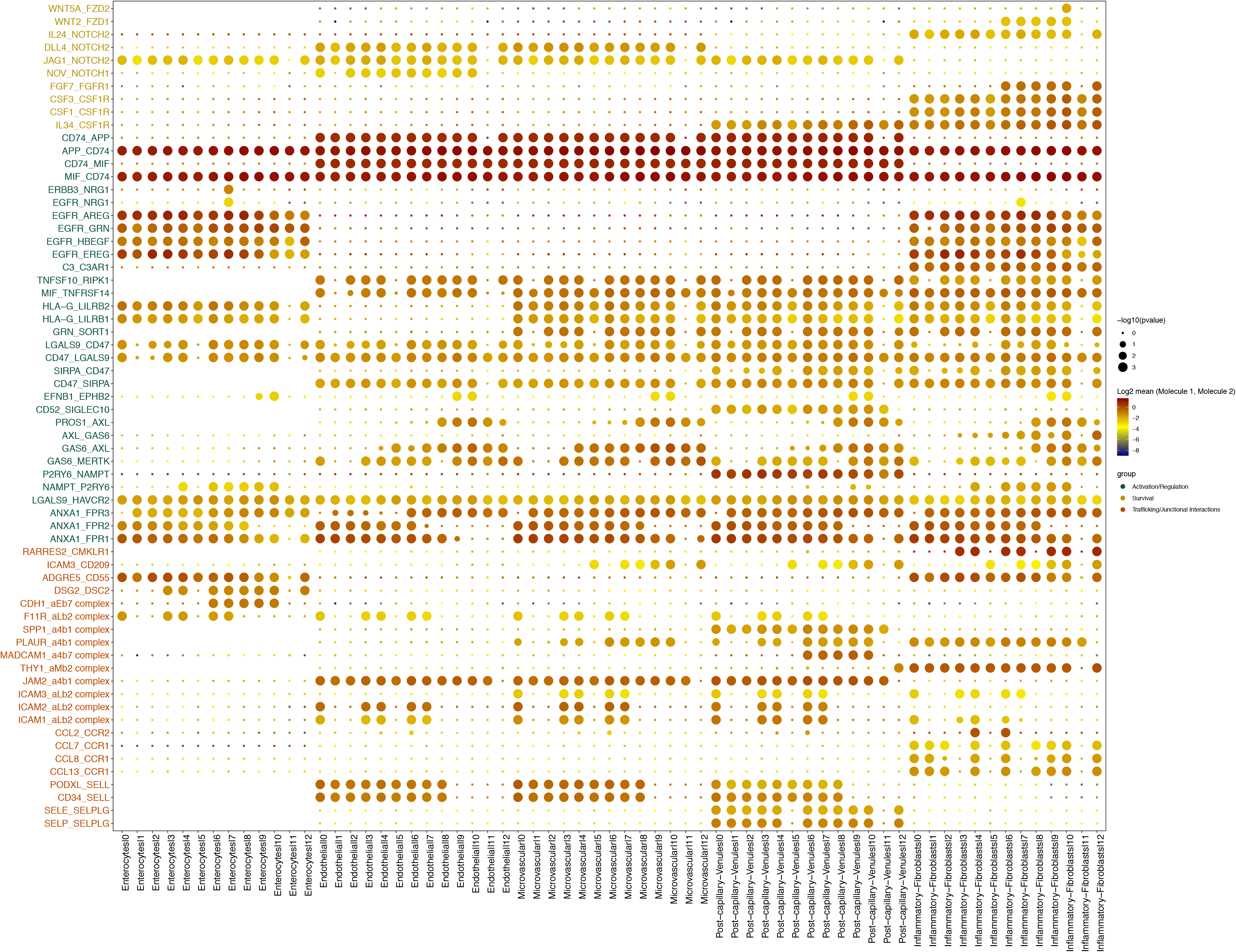
Dot plot of interactions between LpM, epithelial cells and stromal cells from colon. Rows represents ligand-receptor pairs and columns defines cell-cell interaction pairs.

**Figure S11:**
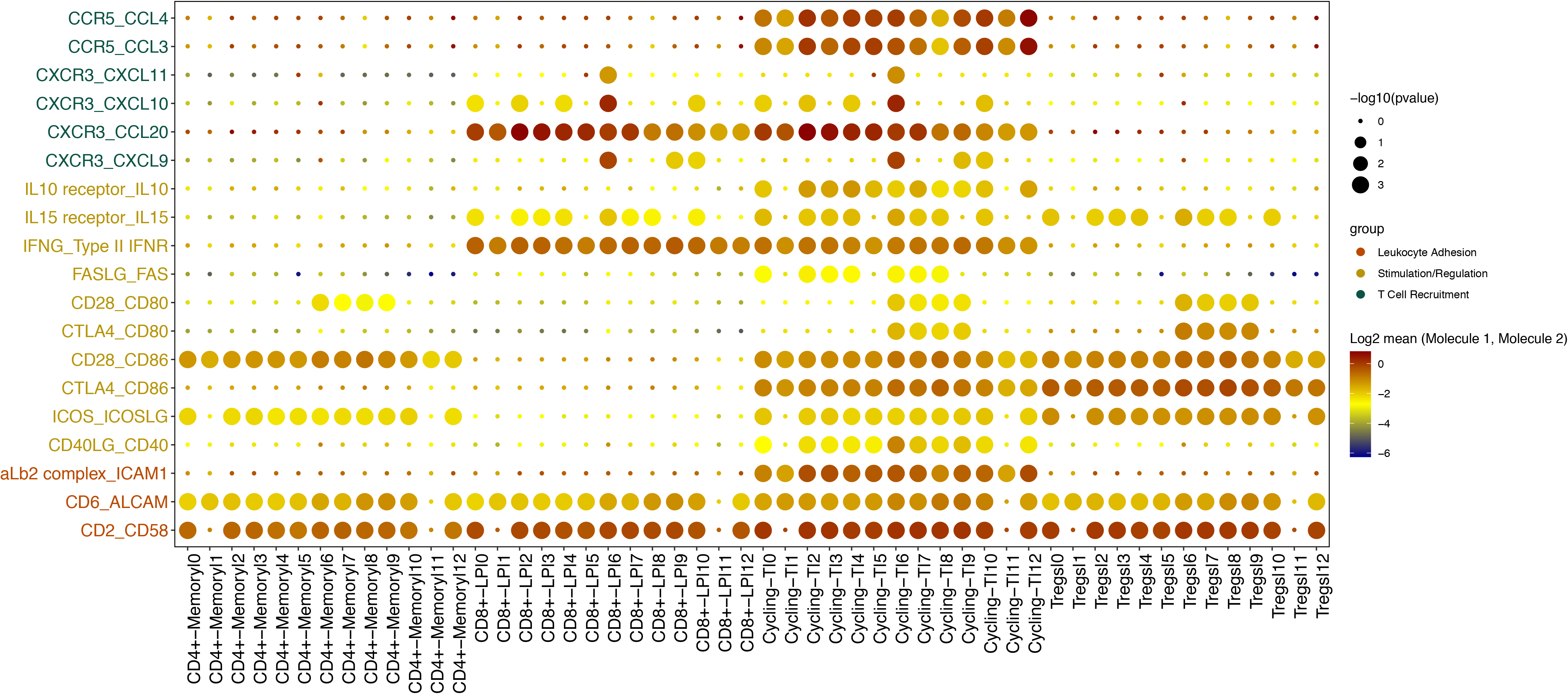
Dot plot of interactions between LpM and immune cells from colon. Rows represents ligand-receptor pairs and columns defines cell-cell interaction pairs.

**Figure S12:**
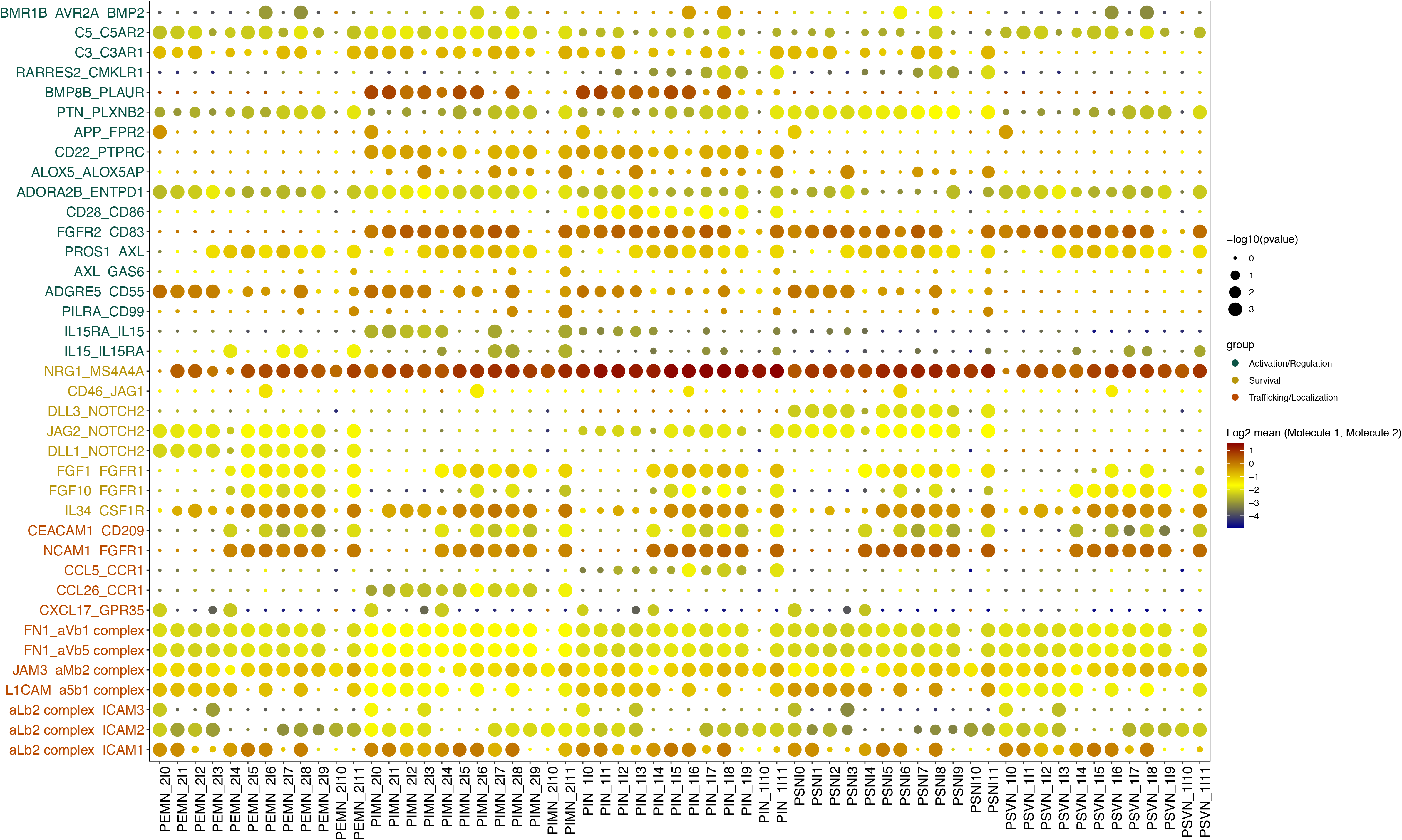
Dot plot of interactions between MM and subtypes of enteric neurons. Rows represents ligand-receptor pairs and columns defines cell-cell interaction pairs.

## References

Arandjelovic, S., and Ravichandran, K.S. (2015). Phagocytosis of apoptotic cells in homeostasis. Nat Immunol 16, 907–917.

Aziz, A., Soucie, E., Sarrazin, S., and Sieweke, M.H. (2009). MafB/c-Maf deficiency enables self-renewal of differentiated functional macrophages. Science 326, 867–871.

Bain, C.C., Bravo-Blas, A., Scott, C.L., Perdiguero, E.G., Geissmann, F., Henri, S., Malissen, B., Osborne, L.C., Artis, D., and Mowat, A.M. (2014). Constant replenishment from circulating monocytes maintains the macrophage pool in the intestine of adult mice. Nat Immunol 15, 929–937.

Barkal, A.A., Brewer, R.E., Markovic, M., Kowarsky, M., Barkal, S.A., Zaro, B.W., Krishnan, V., Hatakeyama, J., Dorigo, O., Barkal, L.J., and Weissman, I.L. (2019). CD24 signalling through macrophage Siglec-10 is a target for cancer immunotherapy. Nature 572, 392–396.

Bleriot, C., Chakarov, S., and Ginhoux, F. (2020). Determinants of Resident Tissue Macrophage Identity and Function. Immunity 52, 957–970.

Bogie, J.F., Mailleux, J., Wouters, E., Jorissen, W., Grajchen, E., Vanmol, J., Wouters, K., Hellings, N., van Horssen, J., Vanmierlo, T., and Hendriks, J.J. (2017). Scavenger receptor collectin placenta 1 is a novel receptor involved in the uptake of myelin by phagocytes. Sci Rep 7, 44794.

Bourdely, P., Anselmi, G., Vaivode, K., Ramos, R.N., Missolo-Koussou, Y., Hidalgo, S., Tosselo, J., Nunez, N., Richer, W., Vincent-Salomon, A., et al. (2020). Transcriptional and Functional Analysis of CD1c(+) Human Dendritic Cells Identifies a CD163(+) Subset Priming CD8(+)CD103(+) T Cells. Immunity 53, 335–352 e338.

Brykczynska, U., Geigges, M., Wiedemann, S.J., Dror, E., Boni-Schnetzler, M., Hess, C., Donath, M.Y., and Paro, R. (2020). Distinct Transcriptional Responses across Tissue-Resident Macrophages to Short-Term and Long-Term Metabolic Challenge. Cell Rep 30, 1627–1643 e1627.

Bujko, A., Atlasy, N., Landsverk, O.J.B., Richter, L., Yaqub, S., Horneland, R., Øyen, O., Aandahl, E.M., Aabakken, L., Stunnenberg, H.G., et al. (2018). Transcriptional and functional profiling defines human small intestinal macrophage subsets. J Exp Med 215, 441–458.

Chakarov, S., Lim, H.Y., Tan, L., Lim, S.Y., See, P., Lum, J., Zhang, X.M., Foo, S., Nakamizo, S., Duan, K., et al. (2019). Two distinct interstitial macrophage populations coexist across tissues in specific subtissular niches. Science 363.

Chen, H.M., van der Touw, W., Wang, Y.S., Kang, K., Mai, S., Zhang, J., Alsina-Beauchamp, D., Duty, J.A., Mungamuri, S.K., Zhang, B., et al. (2018). Blocking immunoinhibitory receptor LILRB2 reprograms tumor-associated myeloid cells and promotes antitumor immunity. J Clin Invest 128, 5647–5662.

Chikina, A.S., Nadalin, F., Maurin, M., San-Roman, M., Thomas-Bonafos, T., Li, X.V., Lameiras, S., Baulande, S., Henri, S., Malissen, B., et al. (2020). Macrophages Maintain Epithelium Integrity by Limiting Fungal Product Absorption. Cell 183, 411–428 e416.

De Schepper, S., Verheijden, S., Aguilera-Lizarraga, J., Viola, M.F., Boesmans, W., Stakenborg, N., Voytyuk, I., Schmidt, I., Boeckx, B., Dierckx de Casterle, I., et al. (2018). Self-Maintaining Gut Macrophages Are Essential for Intestinal Homeostasis. Cell 175, 400–415 e413.

Drokhlyansky, E., Smillie, C.S., Van Wittenberghe, N., Ericsson, M., Griffin, G.K., Eraslan, G., Dionne, D., Cuoco, M.S., Goder-Reiser, M.N., Sharova, T., et al. (2020). The Human and Mouse Enteric Nervous System at Single-Cell Resolution. Cell 182, 1606–1622 e1623.

Earley, A.M., Graves, C.L., and Shiau, C.E. (2018). Critical Role for a Subset of Intestinal Macrophages in Shaping Gut Microbiota in Adult Zebrafish. Cell Rep 25, 424–436.

Efremova, M., Vento-Tormo, M., Teichmann, S.A., and Vento-Tormo, R. (2020). CellPhoneDB: inferring cell-cell communication from combined expression of multi-subunit ligand-receptor complexes. Nat Protoc 15, 1484–1506.

Eguiluz-Gracia, I., Schultz, H.H., Sikkeland, L.I., Danilova, E., Holm, A.M., Pronk, C.J., Agace, W.W., Iversen, M., Andersen, C., Jahnsen, F.L., and Baekkevold, E.S. (2016). Long-term persistence of human donor alveolar macrophages in lung transplant recipients. Thorax 71, 1006–1011.

Fawkner-Corbett, D., Antanaviciute, A., Parikh, K., Jagielowicz, M., Geros, A.S., Gupta, T., Ashley, N., Khamis, D., Fowler, D., Morrissey, E., et al. (2021). Spatiotemporal analysis of human intestinal development at single-cell resolution. Cell 184, 810–826 e823.

Gabanyi, I., Muller, P.A., Feighery, L., Oliveira, T.Y., Costa-Pinto, F.A., and Mucida, D. (2016). Neuro-immune Interactions Drive Tissue Programming in Intestinal Macrophages. Cell 164, 378–391.

Gao, X., Hu, D., Gogol, M., and Li, H. (2019). ClusterMap: compare multiple single cell RNA-Seq datasets across different experimental conditions. Bioinformatics 35, 3038–3045.

Godwin, J.W., Pinto, A.R., and Rosenthal, N.A. (2013). Macrophages are required for adult salamander limb regeneration. Proc Natl Acad Sci U S A 110, 9415–9420.

Guilliams, M., De Kleer, I., Henri, S., Post, S., Vanhoutte, L., De Prijck, S., Deswarte, K., Malissen, B., Hammad, H., and Lambrecht, B.N. (2013). Alveolar macrophages develop from fetal monocytes that differentiate into long-lived cells in the first week of life via GM-CSF. J Exp Med 210, 1977–1992.

Guilliams, M., Thierry, G.R., Bonnardel, J., and Bajenoff, M. (2020). Establishment and Maintenance of the Macrophage Niche. Immunity 52, 434–451.

Hartwig, T., Montinaro, A., von Karstedt, S., Sevko, A., Surinova, S., Chakravarthy, A., Taraborrelli, L., Draber, P., Lafont, E., Arce Vargas, F., et al. (2017). The TRAIL-Induced Cancer Secretome Promotes a Tumor-Supportive Immune Microenvironment via CCR2. Mol Cell 65, 730–742 e735.

Hong, S., Beja-Glasser, V.F., Nfonoyim, B.M., Frouin, A., Li, S., Ramakrishnan, S., Merry, K.M., Shi, Q., Rosenthal, A., Barres, B.A., et al. (2016). Complement and microglia mediate early synapse loss in Alzheimer mouse models. Science 352, 712–716.

Hulsmans, M., Clauss, S., Xiao, L., Aguirre, A.D., King, K.R., Hanley, A., Hucker, W.J., Wulfers, E.M., Seemann, G., Courties, G., et al. (2017). Macrophages Facilitate Electrical Conduction in the Heart. Cell 169, 510–522 e520.

Jahchan, N.S., Mujal, A.M., Pollack, J.L., Binnewies, M., Sriram, V., Reyno, L., and Krummel, M.F. (2019). Tuning the Tumor Myeloid Microenvironment to Fight Cancer. Front Immunol 10, 1611.

Jaitin, D.A., Adlung, L., Thaiss, C.A., Weiner, A., Li, B., Descamps, H., Lundgren, P., Bleriot, C., Liu, Z., Deczkowska, A., et al. (2019). Lipid-Associated Macrophages Control Metabolic Homeostasis in a Trem2-Dependent Manner. Cell 178, 686–698 e614.

Katzenelenbogen, Y., Sheban, F., Yalin, A., Yofe, I., Svetlichnyy, D., Jaitin, D.A., Bornstein, C., Moshe, A., Keren-Shaul, H., Cohen, M., et al. (2020). Coupled scRNA-Seq and Intracellular Protein Activity Reveal an Immunosuppressive Role of TREM2 in Cancer. Cell 182, 872–885 e819.

Lavin, Y., Mortha, A., Rahman, A., and Merad, M. (2015). Regulation of macrophage development and function in peripheral tissues. Nat Rev Immunol 15, 731–744.

Lavin, Y., Winter, D., Blecher-Gonen, R., David, E., Keren-Shaul, H., Merad, M., Jung, S., and Amit, I. (2014). Tissue-resident macrophage enhancer landscapes are shaped by the local microenvironment. Cell 159, 1312–1326.

Li, Z., Li, Y., Gao, J., Fu, Y., Hua, P., Jing, Y., Cai, M., Wang, H., and Tong, T. (2021). The role of CD47-SIRPalpha immune checkpoint in tumor immune evasion and innate immunotherapy. Life Sci 273, 119150.

Lim, H.Y., Lim, S.Y., Tan, C.K., Thiam, C.H., Goh, C.C., Carbajo, D., Chew, S.H.S., See, P., Chakarov, S., Wang, X.N., et al. (2018). Hyaluronan Receptor LYVE-1-Expressing Macrophages Maintain Arterial Tone through Hyaluronan-Mediated Regulation of Smooth Muscle Cell Collagen. Immunity 49, 326–341 e327.

Luoma, A.M., Suo, S., Williams, H.L., Sharova, T., Sullivan, K., Manos, M., Bowling, P., Hodi, F.S., Rahma, O., Sullivan, R.J., et al. (2020). Molecular Pathways of Colon Inflammation Induced by Cancer Immunotherapy. Cell 182, 655–671 e622.

Martin, J.C., Chang, C., Boschetti, G., Ungaro, R., Giri, M., Grout, J.A., Gettler, K., Chuang, L.S., Nayar, S., Greenstein, A.J., et al. (2019). Single-Cell Analysis of Crohn’s Disease Lesions Identifies a Pathogenic Cellular Module Associated with Resistance to Anti-TNF Therapy. Cell 178, 1493–1508.e1420.

Matheis, F., Muller, P.A., Graves, C.L., Gabanyi, I., Kerner, Z.J., Costa-Borges, D., Ahrends, T., Rosenstiel, P., and Mucida, D. (2020). Adrenergic Signaling in Muscularis Macrophages Limits Infection-Induced Neuronal Loss. Cell 180, 64–78 e16.

Muller, P.A., Koscso, B., Rajani, G.M., Stevanovic, K., Berres, M.L., Hashimoto, D., Mortha, A., Leboeuf, M., Li, X.M., Mucida, D., et al. (2014). Crosstalk between Muscularis Macrophages and Enteric Neurons Regulates Gastrointestinal Motility. Cell 158, 1210.

Myers, K.V., Amend, S.R., and Pienta, K.J. (2019). Targeting Tyro3, Axl and MerTK (TAM receptors): implications for macrophages in the tumor microenvironment. Mol Cancer 18, 94.

Ocana-Guzman, R., Torre-Bouscoulet, L., and Sada-Ovalle, I. (2016). TIM-3 Regulates Distinct Functions in Macrophages. Front Immunol 7, 229.

Okabe, Y., and Medzhitov, R. (2014). Tissue-specific signals control reversible program of localization and functional polarization of macrophages. Cell 157, 832–844.

Patel, A.A., Ginhoux, F., and Yona, S. (2021). Monocytes, macrophages, dendritic cells and neutrophils: an update on lifespan kinetics in health and disease. Immunology.

Qu, Y., Wen, J., Thomas, G., Yang, W., Prior, W., He, W., Sundar, P., Wang, X., Potluri, S., and Salek-Ardakani, S. (2020). Baseline Frequency of Inflammatory Cxcl9-Expressing Tumor-Associated Macrophages Predicts Response to Avelumab Treatment. Cell Rep 32, 108115.

Ramachandran, P., Dobie, R., Wilson-Kanamori, J.R., Dora, E.F., Henderson, B.E.P., Luu, N.T., Portman, J.R., Matchett, K.P., Brice, M., Marwick, J.A., et al. (2019). Resolving the fibrotic niche of human liver cirrhosis at single-cell level. Nature 575, 512–518.

Sharma, A., Paranjape, A.N., Rangarajan, A., and Dighe, R.R. (2012). A monoclonal antibody against human Notch1 ligand-binding domain depletes subpopulation of putative breast cancer stem-like cells. Mol Cancer Ther 11, 77–86.

Sharma, A., Seow, J.J.W., Dutertre, C.A., Pai, R., Bleriot, C., Mishra, A., Wong, R.M.M., Singh, G.S.N., Sudhagar, S., Khalilnezhad, S., et al. (2020). Onco-fetal Reprogramming of Endothelial Cells Drives Immunosuppressive Macrophages in Hepatocellular Carcinoma. Cell 183, 377–394 e321.

Shaw, T.N., Houston, S.A., Wemyss, K., Bridgeman, H.M., Barbera, T.A., Zangerle-Murray, T., Strangward, P., Ridley, A.J.L., Wang, P., Tamoutounour, S., et al. (2018). Tissue-resident macrophages in the intestine are long lived and defined by Tim-4 and CD4 expression. J Exp Med 215, 1507–1518.

Smillie, C.S., Biton, M., Ordovas-Montanes, J., Sullivan, K.M., Burgin, G., Graham, D.B., Herbst, R.H., Rogel, N., Slyper, M., Waldman, J., et al. (2019). Intra- and Inter-cellular Rewiring of the Human Colon during Ulcerative Colitis. Cell 178, 714–730.e722.

Stephan, A.H., Barres, B.A., and Stevens, B. (2012). The complement system: an unexpected role in synaptic pruning during development and disease. Annu Rev Neurosci 35, 369–389.

Stuart, T., Butler, A., Hoffman, P., Hafemeister, C., Papalexi, E., Mauck, W.M., 3rd, Hao, Y., Stoeckius, M., Smibert, P., and Satija, R. (2019). Comprehensive Integration of Single-Cell Data. Cell 177, 1888–1902 e1821.

Taylor, V., Welcher, A.A., Program, A.E., and Suter, U. (1995). Epithelial membrane protein-1, peripheral myelin protein 22, and lens membrane protein 20 define a novel gene family. J Biol Chem 270, 28824–28833.

Trapnell, C., Cacchiarelli, D., Grimsby, J., Pokharel, P., Li, S., Morse, M., Lennon, N.J., Livak, K.J., Mikkelsen, T.S., and Rinn, J.L. (2014). The dynamics and regulators of cell fate decisions are revealed by pseudotemporal ordering of single cells. Nat Biotechnol 32, 381–386.

Van de Sande, B., Flerin, C., Davie, K., De Waegeneer, M., Hulselmans, G., Aibar, S., Seurinck, R., Saelens, W., Cannoodt, R., Rouchon, Q., et al. (2020). A scalable SCENIC workflow for single-cell gene regulatory network analysis. Nat Protoc 15, 2247–2276.

Wang, W., Marinis, J.M., Beal, A.M., Savadkar, S., Wu, Y., Khan, M., Taunk, P.S., Wu, N., Su, W., Wu, J., et al. (2018). RIP1 Kinase Drives Macrophage-Mediated Adaptive Immune Tolerance in Pancreatic Cancer. Cancer Cell 34, 757–774 e757.

Zhang, L., Li, Z., Skrzypczynska, K.M., Fang, Q., Zhang, W., O’Brien, S.A., He, Y., Wang, L., Zhang, Q., Kim, A., et al. (2020). Single-Cell Analyses Inform Mechanisms of Myeloid-Targeted Therapies in Colon Cancer. Cell 181, 442–459 e429.

